# Root-to-shoot iron partitioning in Arabidopsis requires IRON-REGULATED TRANSPORTER1 (IRT1)

**DOI:** 10.1101/2021.02.08.430285

**Authors:** Julia Quintana, María I. Bernal, Marleen Scholle, Heike Holländer-Czytko, Nguyen Nga, Markus Piotrowski, David G. Mendoza-Cózatl, Michael J. Haydon, Ute Krämer

**Affiliations:** Faculty of Biology and Biotechnology, Ruhr University Bochum, 44801 Bochum, Germany; Department of Plant Nutrition, Estación Experimental de Aula Dei-CSIC, 50059 Zaragoza, Spain; Division of Plant Sciences, MU-Columbia, Columbia, MO 65211-7310, USA; School of BioSciences, University of Melbourne, Melbourne, Australia

## Abstract

IRON-REGULATED TRANSPORTER1 (IRT1) is the root high-affinity ferrous iron uptake system and indispensable for the completion of the life cycle of *Arabidopsis thaliana* without vigorous iron (Fe) supplementation. Here we provide evidence supporting a second role of IRT1 in root-to-shoot mobilization of Fe. We show that the *irt1*-2 (*pam42*) mutant over-accumulates Fe in roots, most prominently in the cortex of the differentiation zone, when compared to the wild type. Shoots of *irt1*-2 are severely Fe-deficient according to Fe content and marker transcripts, as expected. We generated *irt1*-2 lines producing IRT1 mutant variants carrying single amino-acid substitutions of key residues in transmembrane helices IV and V, Ser_206_ and His_232_, which are required for transport activity in yeast. In the transgenic Arabidopsis lines, short-term root Fe uptake rates and secondary substrate Mn accumulation resemble those of *irt1*-2, suggesting that these plants remain incapable of IRT1-mediated root Fe uptake. Yet, IRT1_S206A_ partially complements rosette dwarfing and leaf chlorosis, as well as root-to-shoot Fe partitioning and gene expression defects of *irt1*-2, all of which are fully complemented by wild-type IRT1. Taken together, these results suggest a function for IRT1 in root-to-shoot Fe partitioning that does not require Fe transport activity of IRT1. Among the genes of which transcript levels are partially dependent on IRT1, we identify *MYB DOMAIN PROTEIN10*, *MYB DOMAIN PROTEIN72* and *NICOTIANAMINE SYNTHASE4* as candidates for effecting IRT1-dependent Fe mobilization in roots. Understanding the biological functions of IRT1 will help to improve iron nutrition and the nutritional quality of agricultural crops.

## INTRODUCTION

The evolution of core components of electron transport chains and central metabolic pathways predated the Great Oxidation Event, which was accompanied by a dramatic decrease in the bioavailability of iron (Fe) approximately 2 Ga ago (Alberts et al., 2002). In all living organisms, the nutritional requirement for the micronutrient Fe is thus generally far higher than bioavailable concentrations in today’s biosphere. Fe is the third most limiting mineral nutrient in agriculture (Zuo and Zhang, 2011), and one-third of the World’s population suffer from Fe-deficiency anemia (Lopez et al., 2016). As primary producers, land plants possess particularly effective mechanisms for mining Fe from the Earth’s crust, and bio-available Fe in plant biomass contributes to sustaining ecosystems and human nutrition.

IRON REGULATED TRANSPORTER1 (IRT1) is the *bona fide* high-affinity root Fe^2+^ uptake system and of central importance in plants pursuing strategy I of iron acquisition, including *Arabidopsis thaliana* (Henriques et al., 2002; Varotto et al., 2002; Vert et al., 2002; Eide et al., 1996; Connolly et al., 2002; Kobayashi and Nishizawa, 2012). Arabidopsis *irt1* mutants have severely chlorotic leaves, are strongly impaired in growth and die before reaching the reproductive stage unless supplemented with Fe chelates. The IRT1 protein forms a complex with the FERRIC REDUCTION OXIDASE2 (FRO2) and AUTOINHIBITED PLASMA MEMBRANE H^+^-ATPase2 (AHA2) for Fe^(+III)^ solubilization and Fe^2+^ uptake into epidermal cells in the root differentiation zone (Martín-Barranco et al., 2020). Besides IRT1, the high-affinity Mn transporter NATURAL RESISTANCE-ASSOCIATED MACROPHAGE PROTEIN1 (NRAMP1) is thought to act as a secondary low-affinity Fe^2+^ uptake system (Cailliatte et al., 2010; Castaings et al., 2016).

In Fe-deficient plants, increased transcription of *IRT1*, *FRO2* and *AHA2* is thought to be controlled by the basic HELIX-LOOP-HELIX (bHLH) transcription factor FE-DEFICIENCY INDUCED TRANSCRIPTION FACTOR (FIT, bHLH29) (Colangelo and Guerinot, 2004; Jakoby et al., 2004; Bauer et al., 2007; Schwarz and Bauer, 2020). UPSTREAM REGULATOR of IRT1 (URI)/bHLH121, in turn, mediates the transcriptional activation of *FIT* under Fe deficiency (Gao et al., 2019a; Kim et al., 2019; Gao et al., 2020b; Lei et al., 2020). Low Fe availability induces URI phosphorylation, which activates the formation heterodimers of URI with bHLH34/104 or bHLH115 (ILR3) (Kim et al., 2019; Gao et al., 2020a). These heterodimers then bind to the promoters of several Fe deficiency-activated regulators of Fe homeostasis in roots such as the *MYB DOMAIN PROTEIN10* (*MYB10*)/*MYB DOMAIN PROTEIN72* (*MYB72*) and *bHLH38/39/100/101*. Notably, the interaction of transcription factors bHLH38/39/100/101 with FIT is necessary for the FIT-dependent activation of *IRT1, FRO2* and *AHA2* (Wang et al., 2007; Sivitz et al., 2012). When a physiologically sufficient level of Fe is restored, FIT is polyubiquitinated by BRUTUS-LIKE1/2 (BTSL1/2) and targeted for degradation in the vacuole (Sivitz et al., 2011; Rodríguez-Celma et al., 2019).

IRT1 appears to undergo continuous cycling between the endosomal compartment and the plasma membrane where IRT1 is active (Barberon et al., 2011; Barberon et al., 2014; Ivanov et al., 2014). The monoubiquitination of IRT1 triggers its trafficking from the plasma membrane to the early endosome, which can constitute the first step towards the degradation of IRT1 in the vacuole (Kerkeb et al., 2008; Barberon et al., 2011). Monoubiquitination, internalization and degradation of IRT1 are strongly impaired in synthetic IRT1 variants carrying mutations of both lysine residues K154 and K179 in the cytosolic loop between transmembrane helices III and IV (Kerkeb et al., 2008; Barberon et al., 2011). SORTING NEXIN1 (SNX1) in the early endosome and the phosphatidylinositol-3-phosphate binding protein FYVE-DOMAIN PROTEIN1 (FYVE1) in the late endosome contribute to the cellular sorting of IRT1, including its delivery to the plasma membrane that counteracts the degradation of endocytosed IRT1 protein (Barberon et al., 2014; Ivanov et al., 2014).

IRT1 lacks specificity for its primary substrate Fe^2+^, so that it can inadvertently transport other cations present in the soil solution such as Mn^2+^, Cd^2+^, Zn^2+^ and Co^2+^ (Eide et al., 1996; Rogers et al., 2000; Vert et al., 2002; Connolly et al., 2002). For example, IRT1 was concluded to be the major route for the intake of non-essential hazardous Cd^2+^ cations into Arabidopsis and likely also into many food crops (Vert et al., 2002). To counteract the accumulation of toxic metals under Fe deficiency, excessive cytosolic levels of secondary-substrate metal cations are sensed by binding to a histidine-rich sequence motif in the cytosolic loop located between the transmembrane domains III and IV of IRT1 (Dubeaux et al., 2018). This leads to the phosphorylation of serine and threonine residues in the same cytosolic loop of IRT1 by CBL-INTERACTING PROTEIN KINASE23 (CIPK23), thus triggering the extension of monoubiquitination into K63-polyubiquitination mediated by IRT1 DEGRADATION FACTOR1 (IDF1), followed by the vacuolar degradation of IRT1 (Dubeaux et al., 2018; Shin et al., 2013). FRO2 and AHA2 can also be ubiquitinated, but – different from IRT1 – their post-translational modification is independent of non-target secondary IRT1 substrate metal cations (Martín-Barranco et al., 2020).

Besides its localization at the root surface for Fe^2+^ cation uptake by the plant, IRT1 was also detected in the root vasculature (Barberon et al., 2011; Marquès-Bueno et al., 2016). Only the combined expression of *IRT1* in both, trichoblasts and phloem companion cells, complemented the dwarfed growth, leaf chlorosis, and infertility of the *irt1* mutant cultivated in soil (Marquès-Bueno et al., 2016). Moreover, the *IRT1* mRNA was reported to move from the shoot into the root, suggesting its cell-to-cell mobility (Thieme et al., 2015). The function of *IRT1* expression in phloem companion cells has remained elusive to date.

Here we provide evidence for a secondary role of IRT1 in root-to-shoot Fe partitioning, independent of its transmembrane Fe^2+^ transport activity. Our results show an over-accumulation of Fe in roots of the *irt1* mutant, whereas *irt1* shoots are severely Fe-deficient in comparison to wild type plants. We then transformed the *irt1* mutant with *IRT1* constructs encoding transport-inactive mutant protein variants IRT1_S206A_ and IRT1_H232A_. Short-term root Fe uptake rates and accumulation of the secondary IRT1 substrate Mn are indistinguishable from *irt1* in these lines, as expected. Nevertheless, IRT1_S206A_ partially rescues rosette dwarfing and leaf chlorosis, as well as alterations in root-to-shoot Fe mobilization and Fe status marker transcript levels, of *irt1*. Using transcriptomics, we further identify *MYB10*, *MYB72* and their target of transcriptional activation *NICOTIANAMINE SYNTHASE4 (NAS4)* as candidates for downstream involvement in IRT1-dependent root-to-shoot Fe partitioning, among other genes.

## RESULTS

### Root-to-shoot iron partitioning is altered in the *irt1* mutant

To examine how Fe concentrations are altered in roots and shoots of the *irt1* mutant, we compared wild-type (WT) and *irt1*-2 seedlings (*pam42*, termed *irt1* below). Seedlings were cultivated under Fe-deficient (− Fe, no added Fe) and Fe-sufficient (+ Fe, 10 μM FeHBED, control) conditions for 10 d, subsequent to a 10-d pre-cultivation period on Fe-sufficient standard medium (Fig. 1A). Upon cultivation in control conditions, shoot Fe concentrations were about one-third lower in *irt1* seedlings than in the WT (Fig. 1B). On Fe-deficient media, Fe concentrations were equivalent in shoots of *irt1* and WT, a known consequence of growth limitation by Fe deficiency (Fig. 1B, Supplemental Fig. S1A and B) (Baxter et al., 2008). In roots, Fe concentrations were about twice as high in *irt1* than in the WT when cultivated under control conditions (Fig. 1C). A similar trend was observed upon cultivation in Fe-deficient media, but the difference between *irt1* and WT was not statistically significant (Fig. 1C). Thus, shoot Fe concentrations were in agreement with expectations, but root Fe concentrations were contrary to expectations, based on the role of IRT1 as the primary root Fe uptake system of Arabidopsis. Congruent with these observations, root-to-shoot Fe concentration ratios were invariably close to unity in WT, but they were strongly shifted towards root Fe accumulation in *irt1* on both control and Fe-deficient media (Fig. 1D). Accounting for condition- and genotype-dependent differences in the degree of Fe limitation of growth (Supplemental Fig. S1A and S1B), total Fe contents (Supplemental Fig. S1C and D) were also in agreement. Taken together, these results suggest a substantial reduction in the partitioning of Fe from roots to shoots in *irt1* mutant plants, in addition to their previously described well-known iron uptake defect (Henriques et al., 2002; Vert et al., 2002). In order to address where over-accumulation of Fe occurs in the roots of *irt1* seedlings, we performed histochemical detection of non-heme Fe^(+II, +III)^ in roots after desorbing apoplastically bound Fe. We confirmed that roots of the constitutively iron-deficient *frd3-7* mutant over-accumulate Fe in the stele, as previously reported (Green and Rogers, 2004). Whole-mount Perls staining of roots of seedlings grown on standard media was in agreement with elevated Fe levels in *irt1* compared to the WT, in which we did not observe any insoluble precipitates of Prussian blue (Fig. 2). In *irt1* roots, the Perls stain was localized in the two sub-epidermal cell layers of the differentiation zone, most prominently in the cortex, and not visible in the stele, different from the *frd3-7* mutant. This observation is consistent with impaired Fe movement into the stele of *irt1*, and it cannot be explained by the loss of the cellular Fe import function of IRT1, which is primarily localized in root epidermal cells (Vert et al., 2002; Marquès-Bueno et al., 2016).

**Figure 1.**
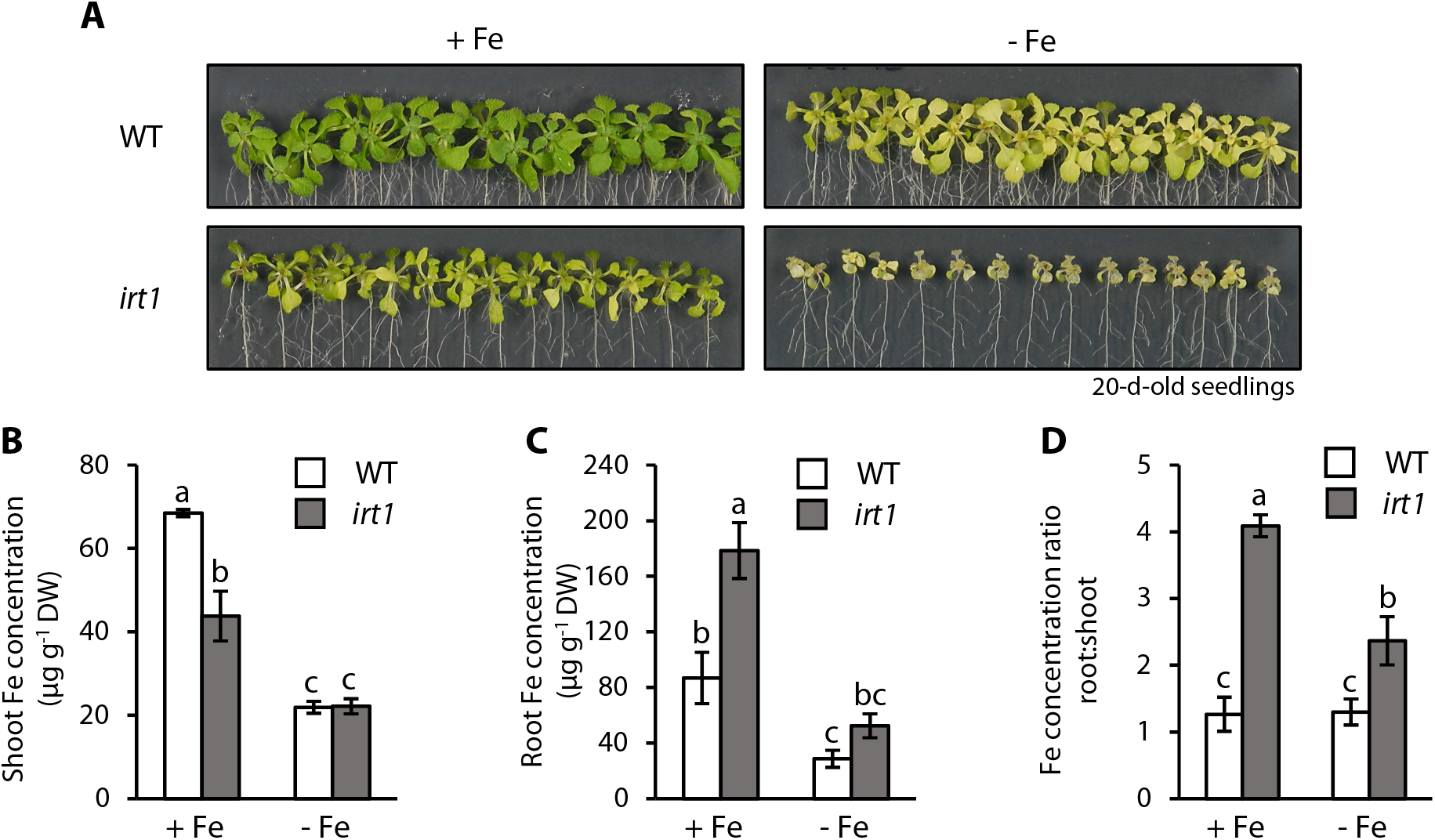
The *irt1* mutant overaccumulates Fe in the root whereas the shoot is Fe-limited. **(A)** Photographs of 20-d-old WT and *irt1* seedlings grown in Fe-sufficient (+ Fe, 10 μM FeHBED) and Fe-deficient (− Fe, 0 μM FeHBED) agar-solidified 0.25x modified Hoagland’s medium (EDTA-washed agar) for 10 d, subsequent to an initial cultivation in standard medium (5 μM FeHBED, unwashed agar) for 10 d, on vertically oriented petri plates. **(B - D)** Shoot **(B)** and root **(C)** Fe concentrations and root:shoot Fe concentration ratio **(D)** in 20-d-old WT and *irt1* seedlings. Seedlings were grown as in **(A)**. Bars represent arithmetic mean ± SD (*n* = 3 pools of tissue, each from 15 (WT) or 30 (*irt1*) seedlings, with 15 seedlings cultivated per plate). Distinct letters indicate statistically significant differences (*P* < 0.05, two-way ANOVA followed by Tukey’s HSD test). Data show one experiment representative of three independent experiments. DW: dry biomass.

**Figure 2.**
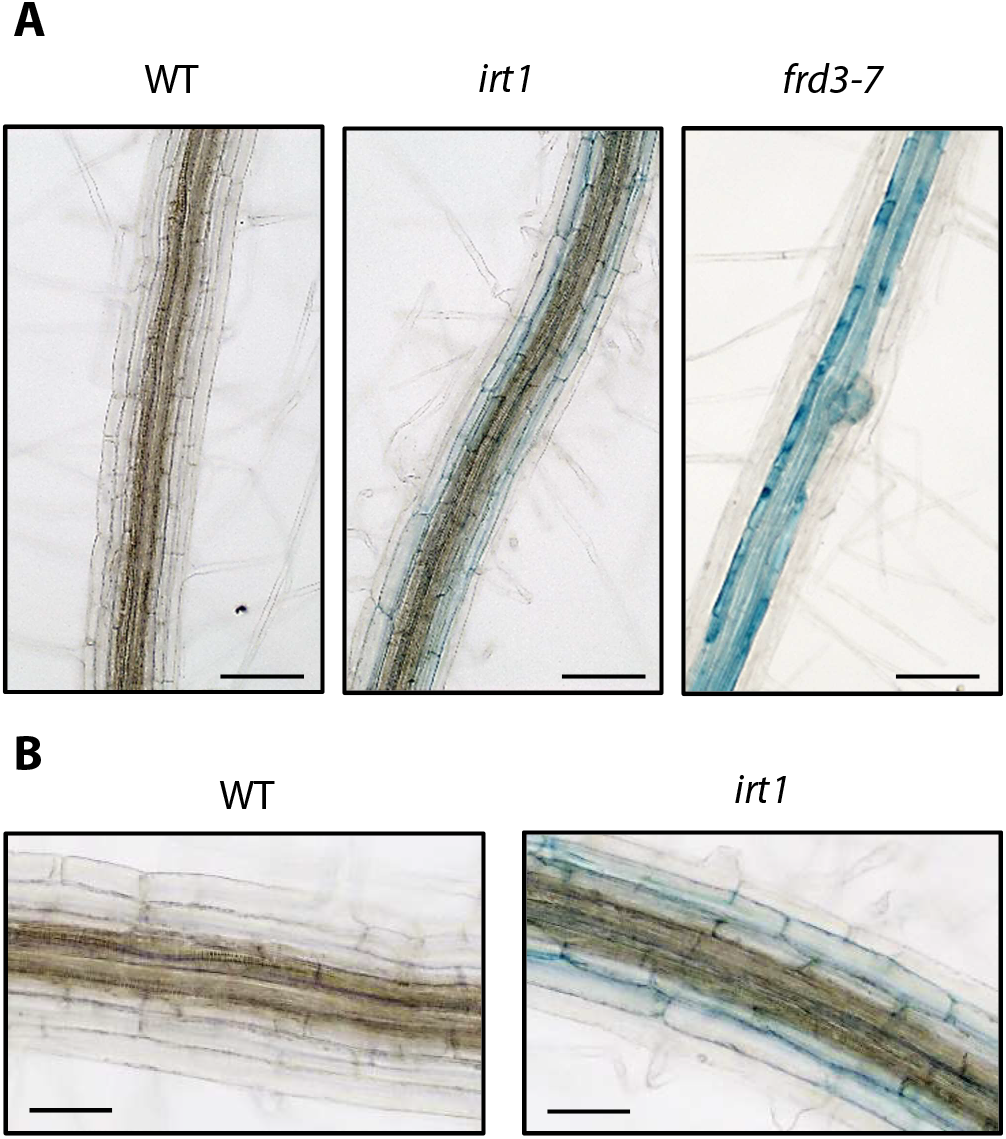
Fe accumulates in sub-epidermal cell layers of *irt1* roots. **(A)** Histochemical detection of labile Fe^(+II, +III)^ by Perls stain in the root differentiation zone of 7-d-old WT, *irt1* and *frd3-7* seedlings grown on agar-solidified 0.25x modified Hoagland’s medium (5 μM FeHBED). **(B)** Magnified image (20x) of WT and *irt1* roots shown in **(A)**. Photographs are representative of *n* = 10 to 12 roots from two independent experiments, each with seedlings cultivated on three replicate petri plates processed in parallel. Scale bars: 100 μm **(A)**, 50 μm **(B)**.

### IRT1_S206A_ variant partially complements the *irt1* mutant

AtIRT1 unequivocally functions in plant Fe uptake by transporting Fe^2+^ cations across the plasma membrane of root epidermal cells (Vert et al., 2002). To test for any additional, distinct role of IRT1 in Fe homeostasis, we used site-directed mutagenesis to construct variants of *IRT1* (Supplemental Fig. S2) that encode IRT1_S206A_ (serine at position 206 substituted by alanine) and IRT1_H232A_ (histidine at position 232 substituted by alanine). Previous work had shown that both of these IRT1 mutant variants were unable to transport both Fe^2+^ and non-target substrates Zn^2+^, Mn^2+^ and Cd^2+^ in the heterologous yeast system *Saccharomyces cerevisiae* (Rogers et al., 2000). Genomic sequences corresponding to the coding region of *IRT1* variants and WT *IRT1* were placed downstream of the native *IRT1* promoter (1,024 bp) and upstream of the native genomic *IRT1* terminator (310 bp). We obtained Arabidopsis *irt1 IRT1_P_:IRT1_S206A_* (termed *irt1* S206A) and *irt1 IRT1_P_:IRT1_H232A_* (*irt1* H232A) lines capable of producing transport-inactive IRT1 only, as well as complemented lines *irt1 IRT1_P_*:*IRT1* (*irt1* IRT1; Supplemental Table S1). *IRT1* transcripts were at comparable levels in the WT and the two complemented lines, and undetectable in the *irt1* mutant (Supplemental Fig. S3A). For both *irt1* S206A and *irt1* H232A, *IRT1* transcript levels were comparable to the WT in one line and approximately tripled those of the WT in the other line, in - Fe-grown seedlings. By comparison, in + Fe-grown seedlings of all genotypes *IRT1* transcript levels were down to a small fraction of between 5% and 16%, except in the *irt1* H232A lines, in which transcript levels in + Fe remained at between 64% and 89% of those in - Fe. Note that in our transgenic *irt1* lines, *IRT1* transcript levels are influenced by both the genomic environment of the T-DNA and the physiological Fe status of seedlings, some of which possess only a transport-inactive IRT1 protein. In contrast to *IRT1*, maximal *IRT2* transcript levels remained below WT levels in all lines and were less than 6.5% of those of *IRT1* in iron-deficient WT seedlings (Supplemental Fig. S3A). Thus, it appears that the T-DNA insertion in *irt1*-2 (*pam42*) within the transcribed region 96 bp upstream of the translational start codon of *IRT1* and about 2,700 bp downstream of the last exon of *IRT2*, also disrupted the function of *IRT2* (Varotto et al., 2002).

IRT1 protein abundance in roots largely reflected *IRT1* transcript levels (Fig. 3A, see Supplemental Fig. S3A). IRT1 protein levels were comparable to the wild type in *irt1* S206A line 2, and elevated compared to both the complemented lines and the wild type in line 1 (Fig. 3A, Supplemental Fig. S3B). Importantly, IRT1 protein was detectable in roots of WT seedlings cultivated on our Fe-sufficient control media, supporting a role of IRT1 in Fe acquisition also under these cultivation conditions (Supplemental Fig. S3B).

**Figure 3.**
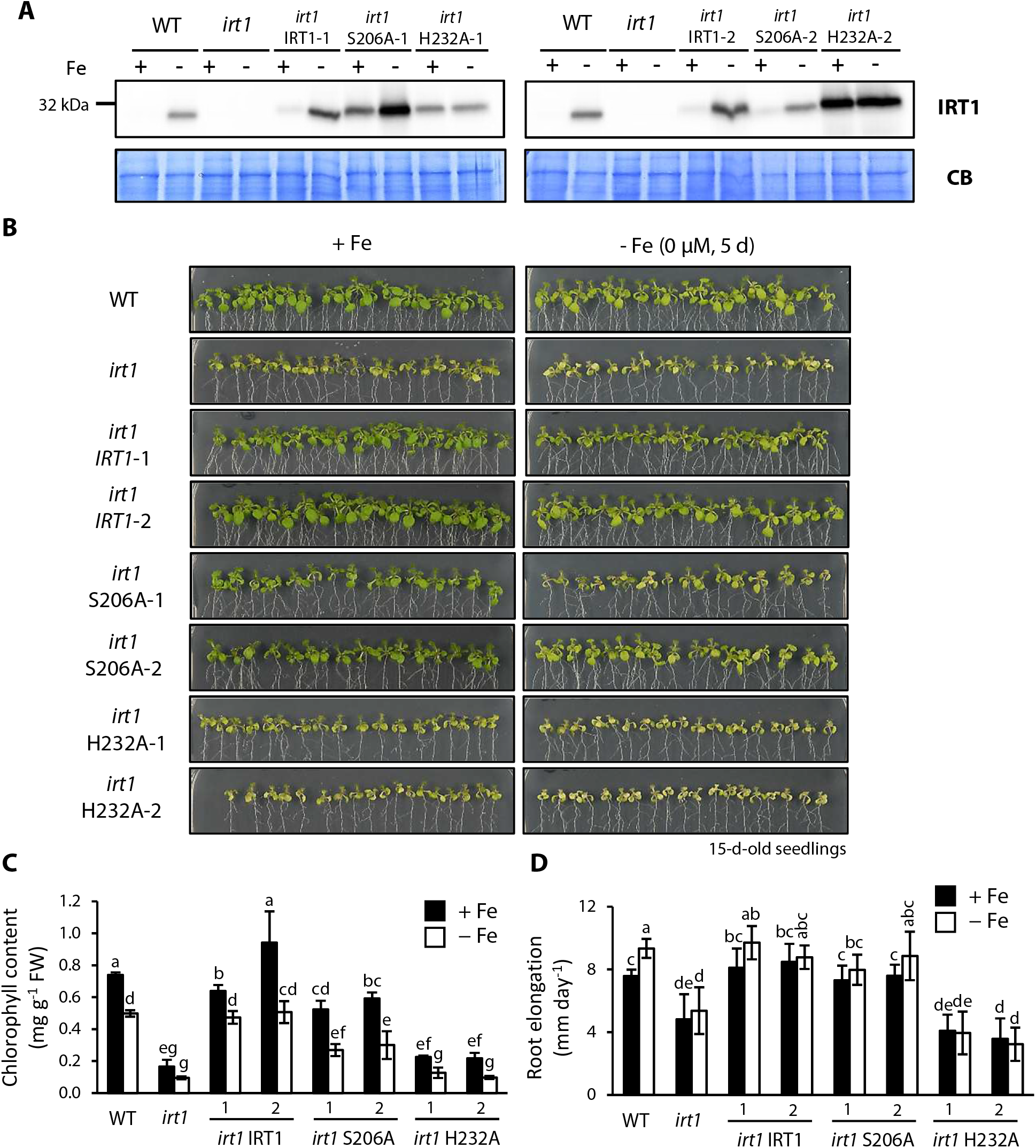
Fe transport-inactive IRT1_S206A_ partially complements *irt1*. **(A - D)** Immunoblot detection of IRT1 protein **(A)**, photographs **(B)**, leaf chlorophyll concentrations **(C)** and mean root elongation over the last 5 d of growth **(D)** in 15-d-old seedlings of WT, *irt1*, *irt1* IRT1, *irt1* S206A and *irt1* H232A. Seedlings were grown on Fe-sufficient (+ Fe, 10 μM FeHBED) and Fe-deficient (− Fe, 0 μM FeHBED) agar-solidified 0.25x modified Hoagland’s medium (EDTA-washed agar) for 5 d, subsequent to an initial cultivation period of 10 d in standard medium (5 μM Fe-HBED; unwashed agar), on vertically oriented petri plates. Total root protein extracts (20 μg) were separated on denaturing gels and blotted onto PVDF membranes, with the Commassie Blue-stained (CB) PVDF membrane shown as loading control **(A)**. Bars represent arithmetic mean ± SD (*n* = 4 pools of 5 to 8 shoots **(C)**; *n* = 8 to 10 roots **(D)**). Distinct letters indicate significant differences (*P* < 0.05) according to two-sample Student’s or Welch *t*-tests upon correction for multiple comparisons **(C)**or two-way ANOVA followed by Tukey’s HSD test **(D)**. Images in **(A)** are representative of three replicate membranes. Data are from two independent experiments, each with two **(C)** or one **(D)** replicate plates.

In the two *irt1* S206A lines, seedling growth and rosette color, leaf chlorophyll concentrations and root growth all suggested a partial complementation of *irt1* mutant phenotypes by comparison to the WT and *irt1*/IRT1 complemented lines (Fig. 3B-D). By contrast, the *irt1* H232A lines resembled the *irt1* mutant throughout. Similarly, soil-grown vegetative *irt1* S206A plants were of a similar size and green color as the WT and *irt1 IRT1* complemented lines, whereas *irt1* H232A lines were indistinguishable from the *irt1* mutant (Supplemental Fig. S4). Rosettes of *irt1* S206A line 2 appeared to be slightly larger and greener than those of line 1 (Fig. 3B, Supplemental Fig. S4). The partial complementation of the *irt1* mutant phenotype through the IRT1_S206A_ variant of an amino acid residue required for transport activity is in agreement with our hypothesis of a secondary biological function of IRT1 in addition to transmembrane Fe^2+^ transport. The IRT1_H232A_ variant of IRT1 was unable to complement the *irt1* mutant phenotype, consistent with a model in which H232 is indispensable for both Fe transport and any additional role in Fe homeostasis carried out by IRT1.

Next we sought to confirm that the *irt1* S206A and *irt1* H232A mutants cannot transport Fe^2+^. In contrast to the WT IRT1 protein, both IRT1_S206A_ and IRT1_H232A_ failed to complement the growth impairment of an iron uptake-defective *S. cerevisiae fet3fet4* mutant on low-iron media (Fig. 4A), as reported previously (Rogers et al., 2000). In order to obtain *in planta* evidence, we quantified ^55^Fe^2+^ incorporation into roots of *ca*. 8-week-old hydroponically cultivated WT, *irt1*, *irt1* IRT1*, irt1* S206A and *irt1* H232A plants after exposure to Fe deficiency for a period of 6 d. Short-term uptake rates of ^55^Fe^2+^ into roots were not decreased in *irt1* by comparison to the WT (Figure 4B, Supplemental Table S2). This result was contrary to expectations, given the lack of the high-affinity root Fe^2+^ uptake system in the *irt1* mutant, and it precluded the complementation testing that we had intended to achieve with this experiment. Importantly, we also detected no differences in root ^55^Fe^2+^ uptake rates between *irt1* S206A and *irt1* H232A, indicating against the differential activation of any secondary Fe uptake systems as the cause for their differing phenotypes (Figure 4B). Altogether, these results suggest that roots of mutants lacking IRT1 transport activity retain significant root Fe^2+^ uptake rates under the employed conditions, which is counter-intuitive but consistent with previously published results including one short-term and one longer-term radiotracer uptake experiment (Henriques et al., 2002; Vert et al., 2002).

**Figure 4.**
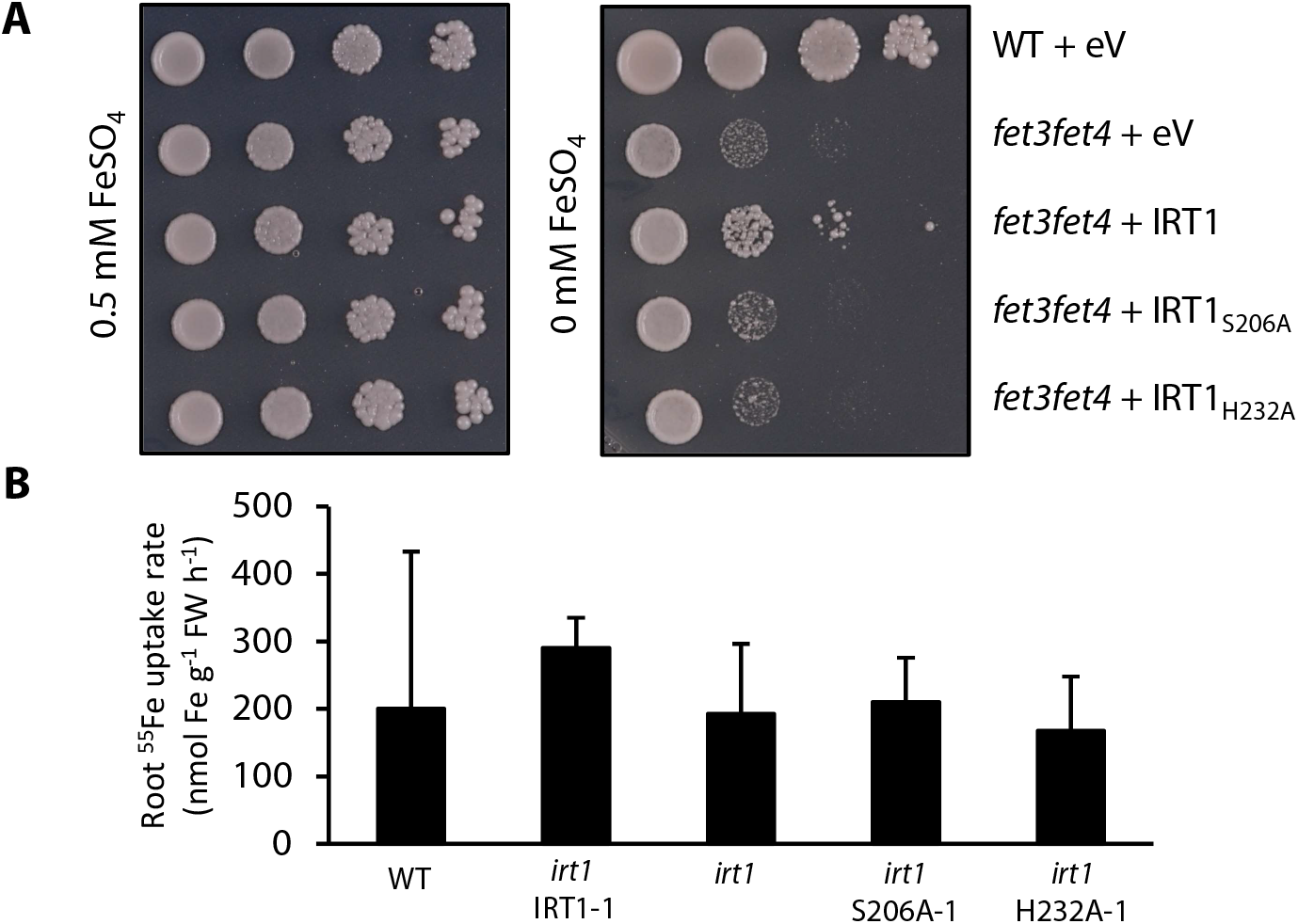
Yeast complementation assay and root ^55^Fe^2+^ uptake rates. **(A)** Growth of WT and the *fet3fet4* Fe transport-defective mutant transformed with a modified pFL61 plasmid as empty vector (eV), or containing the *IRT1*, *IRT1_S206A_* or *IRT1_H232A_* cDNA, respectively. Ten-fold serial dilutions of liquid cultures were spotted on SD (-Ura) medium (pH 5.7) supplemented with 0.5 mM FeSO_4_ or left unamended. **(B)** Uptake rates determined in hydroponically cultivated 7.5 to 8.5-w-old plants over a 13.5-min period. Before the initiation of uptake assays, plants were exposed to Fe deficiency (0 μM FeHBED) for 6 d, subsequent to an initial cultivation period in 1x modified Hoagland‘s solution containing 10 μM FeHBED (WT and *irt1* IRT1) or 50 μM FeHBED (all other genotypes) for 6 to 7 weeks. Bars represent arithmetic mean ± SD (*n* = 4 to 5 plants). Data show one experiment representative of three independent experiments (see Supplemental Table S2).

### The root-to-shoot Fe partitioning defect of *irt1* is partially rescued by the transport-inactive IRT1_S206A_ variant

We observed root Fe over-accumulation in *irt1*, as well as a partial complementation of growth impairment and chlorosis of *irt1* by a transport-inactive IRT1_S206A_ mutant (see Figs. 1C, 2 and 3). Next we addressed root-to-shoot Fe partitioning. Shoot Fe concentrations were significantly higher in *irt1* S206A lines compared to *irt1* following growth in control media, and indistinguishable from *irt1* in *irt1* H232A (Fig. 5A and B). Note that all three of these genotypes are physiologically Fe-deficient in control media because they share the lack of IRT1-mediated high-affinity root Fe uptake. Upon growth on Fe-deficient media, there was little or no difference in shoot Fe concentrations between WT and *irt1* (as noted before, see Fig. 1), indicating that this condition is generally less informative (Figure 5A and B). Fe allocation to shoots corroborated a partial complementation in *irt1* S206A lines of the root-to-shoot Fe partitioning defect of *irt1* (Fig. 5C and D). Shoot Mn concentrations were in agreement with the lack of active IRT1 protein in the roots, because the levels of Mn – as a secondary substrate of IRT1 – were similar in shoots of the *irt1* mutant and the transport-inactive lines (Fig. 5E and F).

**Figure 5.**
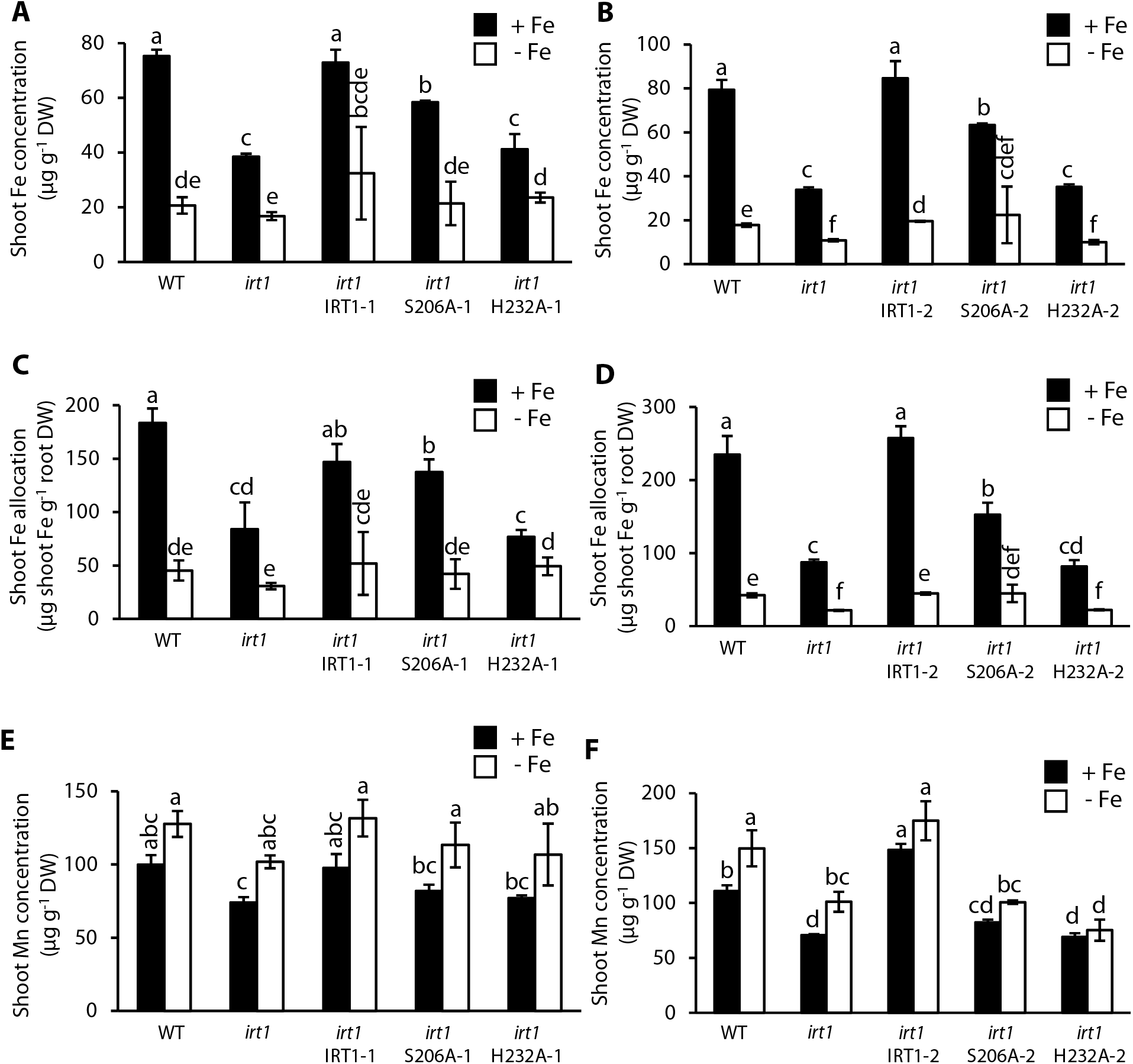
Partial rescue of root-to-shoot Fe allocation in *irt1* S206A. **(A - F)** Shoot Fe concentration **(A and B)**, shoot Fe allocation **(C and D)** and shoot Mn concentrations **(E and F)** in 20-d-old seedlings of WT, *irt1*, *irt1* IRT1, *irt1* S206A and *irt1* H232A grown on Fe-sufficient (+ Fe, 10 μM FeHBED) and Fe-deficient (− Fe, 0 μM FeHBED) agar-solidified 0.25x modified Hoagland’s medium (EDTA-washed agar) for 10 d, subsequent to an initial cultivation period of 10 d in standard medium (5 μM FeHBED; unwashed agar), on vertically oriented petri plates. Bars represent arithmetic mean ± SD (*n* = 3 pools of tissues, each from 15 to 30 seedlings, with 15 seedlings cultivated per plate). Distinct letters indicate significant differences (*P* < 0.05) according to two-sample Student’s *t*-tests upon correction for multiple comparisons **(A - D)** or in two-way ANOVA followed by Tukey’s HSD test **(E and F)**. Data show one experiment representative of two independent experiments.

Root Fe accumulation of *irt1* mutants was restored down to levels resembling the WT in *irt1* S206A seedlings, but not in *irt1* H232A, under control conditions (Supplemental Fig. S5A and B). Note that *irt1 IRT1* lines contained higher *IRT1* transcript and IRT1 protein levels than the WT under + Fe conditions (see Supplemental Fig. S3B), which affects Fe homeostasis, and that *irt1* H232A seedlings of line 2 were very small and chlorotic (see Fig. 3). Total Fe per plant in the *irt1* H232A lines was similar to (*irt1* H232A-1), or even lower (*irt1* H232A-2) than, total Fe in the *irt1* mutant. In *irt1* S206 lines, total Fe was consistently higher than total Fe of *irt1* and the *irt1* H232A lines, and lower than, or similar to, WT and *irt1* IRT1, in control conditions (Supplemental Fig. S5C and D). We propose that increased total Fe levels in the *irt1* S206A lines could be attributed to larger seedling size, a larger root surface area resulting from a longer root in *irt1* S206A than in *irt1* (see Fig. 3), or differing overall Fe homeostasis as a consequence of altered Fe distribution within the plant. In *irt1*, *irt1* S206A and *irt1* H232A, root concentrations of the secondary IRT1 substrate Mn were generally far lower than in both WT and *irt1* IRT1 seedlings, in which they increased under - Fe with increasing IRT1 protein levels, as expected (see Supplemental Fig. S3). Root *NRAMP1* transcript levels were 2 to 3.3-fold elevated in *irt1* and *irt1 H232A* lines, when compared to the WT (Supplemental Fig. S5G). In *irt1* S206A they were lower and remained only slightly, between 1.3- and 1.8-fold, increased compared to *NRAMP1* transcript levels of the WT and *irt1* IRT1 lines. These observations are consistent with IRT1 being the predominant membrane transport protein contributing to root Mn levels under our experimental conditions. In *irt1* S206A-1, root Mn concentrations approximated WT levels under + Fe, although remaining below these on average. The fact that root Mn concentrations did not increase at all on - Fe medium in the presence of increased levels of IRT1_S206A_ protein (see Supplemental Fig. S3) suggested against a direct causal involvement of IRT1_S206A_. Taking these results together, residual IRT1 transport activity or an activation of a secondary Fe uptake system, including *NRAMP1*, cannot explain the partial complementation of shoot Fe levels in *irt1* S206A lines (see Fig. 5A and B). In summary, the root-to-shoot iron partitioning defect of the *irt1* mutant is partially rescued by the transport-inactive IRT1_S206A_, but not IRT1_H232A_.

Ferritins are plastid-localized iron storage proteins, and their levels increase when plants are Fe-replete (Briat et al., 2009). In shoots of WT and *irt1* IRT1 seedlings cultivated on - Fe media for 5 d, *FERRITIN1* (*FER1*) transcript levels were as low as 4.6 to 6.7% of those observed in control media (Fig. 6A). *FER1* expression in *irt1* H232A under control conditions was comparable to that of *irt1* and Fe-deficient WT (Fig. 6A). *FER1* transcript levels in shoots of *irt1* S206A lines were intermediate between WT and *irt1*, in agreement with growth and Fe contents in these lines (Fig. 6A, compare Fig. 5 and Fig. S5). In roots, the differences in *FER1* transcript levels between lines were qualitatively similar, but quantitatively much smaller, in line with root Fe levels (Fig. 6B. compare Fig. 5 and Supplemental Fig. S5). Transcript levels of Fe deficiency marker genes *IRONMAN1* (*IMA1)*, *ZINC INDCED FACILITATOR1* (*ZIF1*), *bHLH39* and *NAS4* in shoots further corroborated a partial complementation of *irt1* by IRT1_S206A_, as evident upon cultivation in control media (Supplemental Fig. S6).

**Figure 6.**
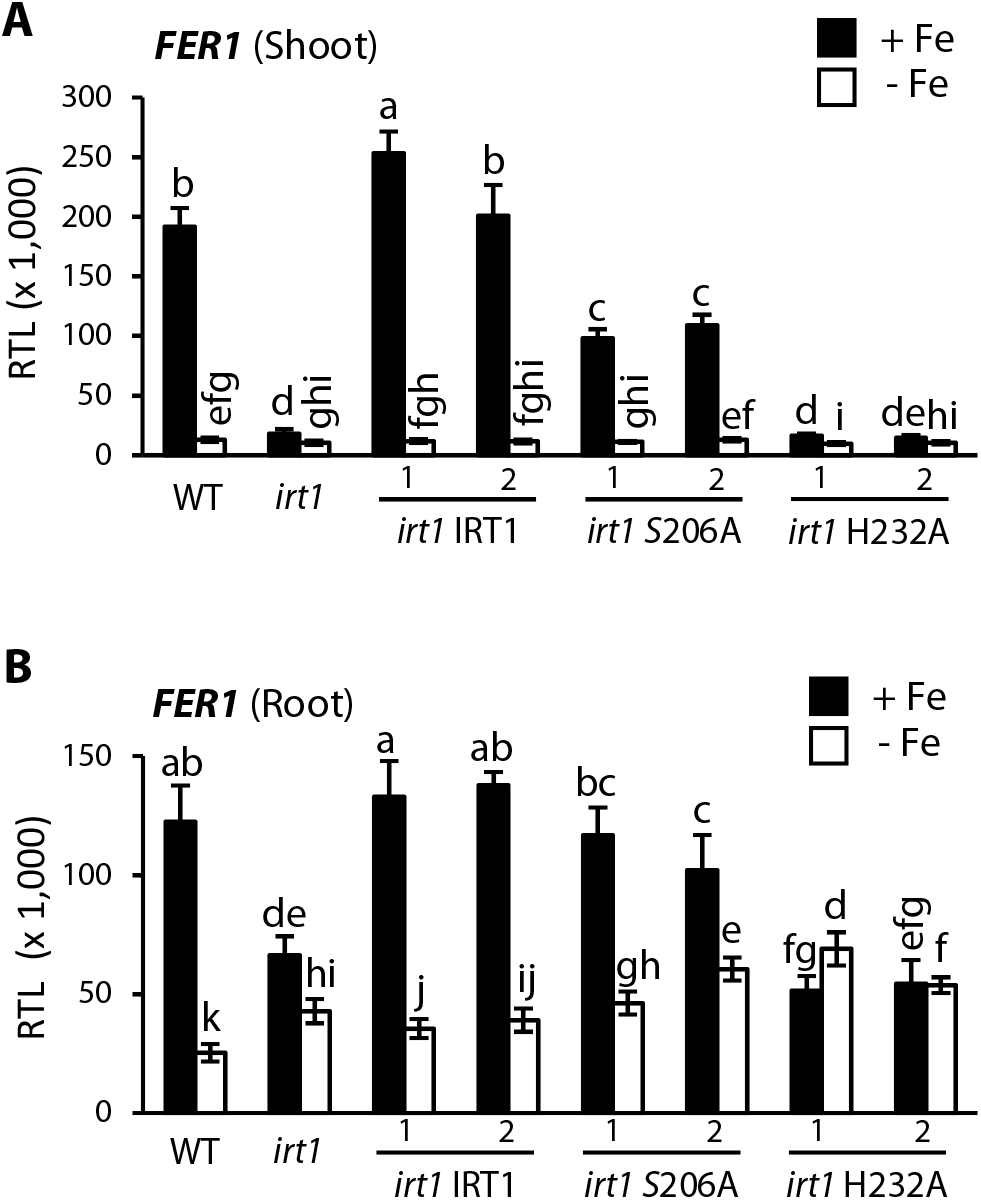
*FER1* transcript levels. **(A - C)** Relative transcript levels (RTL) of *FER1* in shoots **(A)** and roots **(B)** of 15-d-old WT, *irt1*, *irt1* IRT1, *irt1* S206A, *irt1* H232A seedlings. RTL are shown normalized to both *UBQ10* and *EF1α* as constitutively expressed control genes. Seedlings were grown on Fe-sufficient (+ Fe, 10 μM FeHBED) and Fe-deficient (− Fe, 0 μM FeHBED) agar-solidified 0.25x modified Hoagland’s medium (EDTA-washed agar) for 5 d, subsequent to an initial cultivation period of 10 d in standard medium (5 μM FeHBED; unwashed agar), on vertically oriented petri plates. Bars represent arithmetic mean ± SD (*n* = 6 technical replicates on cDNA obtained using tissue pooled from 2 to 3 plates, each with 20 seedlings). Distinct letters indicate significant differences (*P* < 0.05) according to two-sample Student’s or Welch *t*-tests upon correction for multiple comparisons. Data show one experiment representative of three independent experiments.

### How is Fe homeostasis altered in roots of *irt1*?

Next we examined whether Fe status marker genes can additionally provide insights into Fe homeostasis in the *irt1* mutant and thus into possible secondary functions of the IRT1 protein. According to all the marker genes analyzed, the degree of physiological Fe deficiency in shoots of the *irt1* mutant grown in control media was approximately equivalent to, or slightly more severe, than in the WT after 5 d of cultivation on Fe-deficient media (Fig. 6, Supplemental Fig. S6). Note that we quantified biomass and mineral contents of tissues after 10-d treatments in order to avoid experimental noise from low biomass of some samples and the associated sensitivity of our results to contamination, for example from dust particles. Considering roots, *FER1* transcript levels in Fe-deficient WT and *irt1* IRT1 seedlings were 20 to 28% of those observed in Fe-replete media (Fig. 6B). When compared to the WT, *FER1* transcript levels in roots of the *irt1* mutant were reduced to a clearly lesser extent than in shoots after cultivation under control conditions. However, root *FER1* transcript levels did not fully reflect increased root Fe concentrations in *irt1* compared to the WT (compare Fig. 6B with Figs. 1C, 2A). *FER1* transcript levels are highly localized according to the eFP browser in 7-day-old seedlings, with at least 3.5 times higher *FER1* transcript levels in root endodermal and phloem companion cells and at least 2.5 times higher *FER1* transcript levels in root pericycle cells than in root cortex and epidermal cells (Supplemental Fig. S9A) (Brady et al., 2007; Winter et al., 2007). All in all, these results suggest a more Fe-replete status in roots than in shoots of *irt1*, supporting that root-accumulated Fe is at least partly physiologically accessible and thus intracellular in the *irt1* mutant.

### Additional genes of the root Fe-deficiency response are mis-regulated in *irt1*

Next we addressed transcriptome differences between *irt1* and the WT in order to identify candidate genes for mediating decreased root-to-shoot iron mobility in the *irt1* mutant. Between 3 and 5 d of Fe deficiency were previously shown to result in maximal transcriptional responses in roots (Connolly et al., 2002; Connolly et al., 2003; Vert et al., 2003). In our experimental conditions, root surface Ferric Chelate Reductase (FCR) activity in WT seedlings reached a maximum at 5 days of Fe deficiency (Supplemental Fig. S7A). At the time point selected for harvest, FCR activity of the *irt1* mutant was 3.2-fold higher than that of WT under control conditions (+ Fe; Supplemental Fig. S7B). In the WT, FCR activity was 6.6-fold higher under - Fe than under + Fe conditions, and approximately 2-fold higher than in the *irt1* cultivated on + Fe medium (Supplemental Figure S7B).

We conducted microarray-based transcriptomics in 15-day-old WT and *irt1* seedlings grown in Fe-replete media (+ Fe) as well as WT seedlings cultivated in Fe-deficient (− Fe) media for the final 5 d before harvest (see also previous section). A comparison between − Fe and + Fe conditions in WT seedlings identified the transcriptomic Fe deficiency response under our experimental conditions (− Fe *vs.* C (WT)). The comparison between *irt1* and WT cultivated under + Fe conditions identified all symptoms and consequences of the lack of IRT1 (*irt1 vs.* WT (C)). Finally, we compared between *irt1* cultivated under + Fe conditions and WT cultivated under - Fe conditions to identify transcriptome aberrations in the *irt1* mutant (*irt1* (C) *vs.* WT - Fe) (Supplemental Fig. S7C).

In our transcriptomics attempt to identify candidate genes for altered root-to-shoot Fe partitioning in the *irt1* mutant, we expected that candidate gene transcript levels respond to Fe deficiency because IRT1 protein levels also respond to iron deficiency. *IRT1* is expressed in roots, and roots are the primary organ that governs root-to-shoot translocation of mineral nutrients. Among the transcripts of which levels increased more than 2-fold under - Fe compared to control conditions in WT roots, 44% had higher levels in *irt1* than WT under control conditions (Figure 7A, - Fe *vs*. C (WT) ↑ AND *irt1 vs*. WT (C) ↑). Approximately three quarters of these transcripts increased in abundance in *irt1* similarly to the increase in Fe-deficient WT (Supplemental Dataset S2). But for the remaining 26% of these transcripts, this increase was clearly less pronounced than expected from the physiological Fe status of *irt1* plants (Fig. 7A, *irt1 vs*. WT (− Fe) ↓, marked by a single asterisk; see above). This group of 20 genes comprises candidates for an IRT1-dependent function in root-to-shoot partitioning of Fe. The corresponding genes included a number of functionally uncharacterized or poorly characterized genes, as well as previously characterized Fe homeostasis genes including *MYB72*, *IMA1, bHLH101*, *NAS4* and *BETA-GLUCOSIDASE42* (*BGLU42*), for example (Supplemental Dataset S1; note that *bHLH100* and *bHLH38*, which are similar to *bHLH101* in both sequence and biological function, are not represented on the microarray). Of the transcripts that decreased in abundance under - Fe *vs*. C (WT) and also in *irt1 vs*. WT (C), none were under-responsive in *irt1* (Fig. 7B, asterisk).

**Figure 7.**
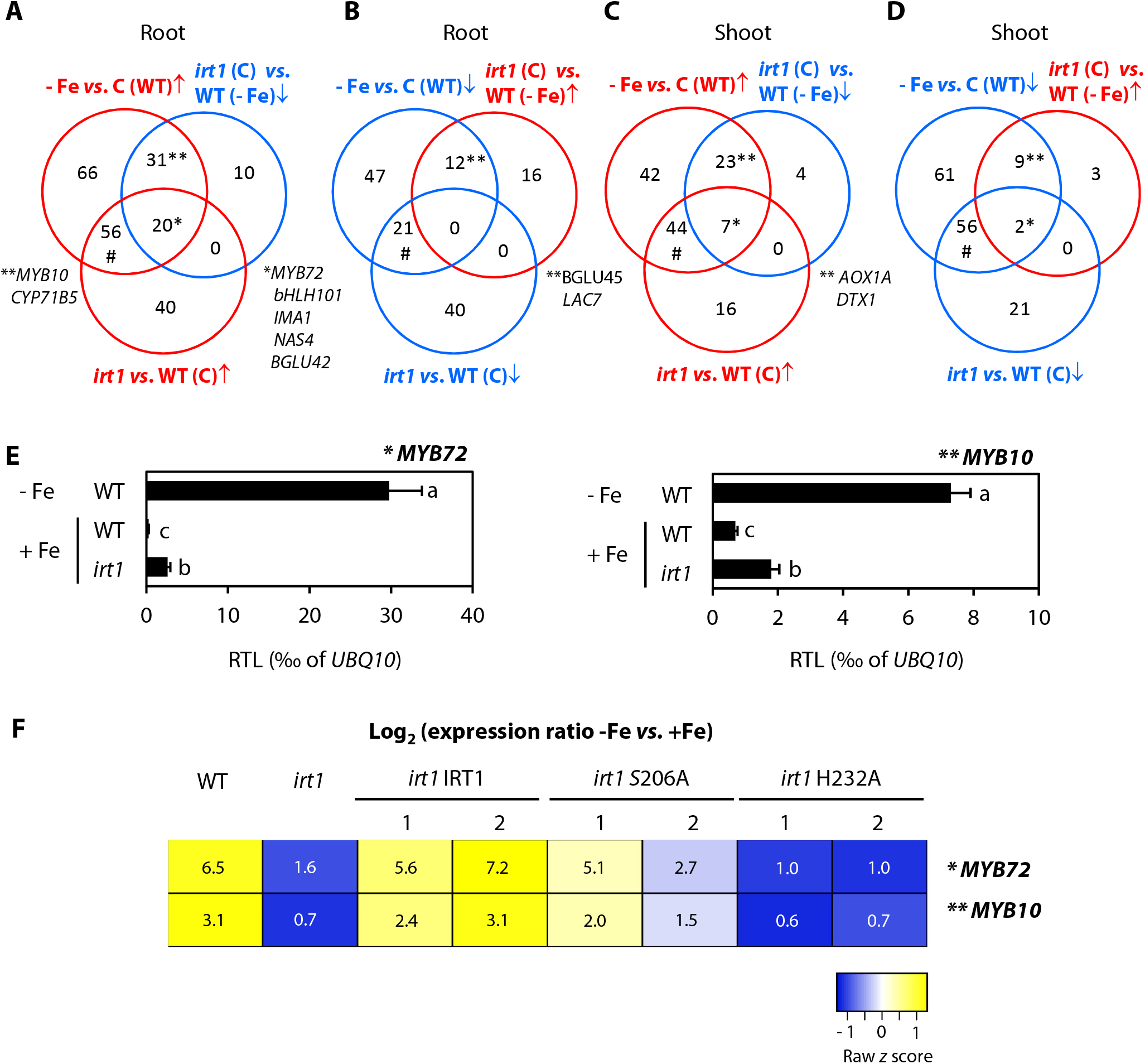
Transcriptome analysis of *irt1* compared to Fe-deficient and -sufficient WT seedlings. **(A - D)** Summary of microarray-based transcriptome data in roots **(A and B)** and shoots **(C and D)**. Shown within Venn diagrams are the numbers of genes with ≥ 2-fold differences in transcript levels in different comparisons (*P* < 0.15). Colors denote responses, (red/blue, higher/lower transcript abundance in one genotype and/or condition compared to the other). Symbols denote Fe deficiency-responsive transcripts that are deregulated (*, **) or not (#) in *irt1*. **(E)** Relative transcript levels (RTL) of *MYB10* and *MYB72* in roots of WT and *irt1* seedlings. RTL are shown normalized to *UBQ10*. **(F)** Heat map of *MYB72* and *MYB10* transcript abundance - Fe *vs*. + Fe in roots of different genotypes as indicated. Numbers are Log_2_ (transcript level ratios, with RTL normalized to both *UBQ10* and *EF1α*), and raw *z*-score for each gene was used to scale the heat map, from three independent experiments. WT and *irt1* seedlings were grown on Fe-sufficient (+ Fe, 10 μM FeHBED) and Fe-deficient (− Fe, 0 μM FeHBED) agar-solidified modified 0.25x Hoagland’s medium (EDTA-washed agar) for 5 d, subsequent to an initial cultivation period of 10 d in standard medium (5 μM FeHBED; unwashed agar), on vertically oriented petri plates **(A - F)**. Bars represent arithmetic mean ± SD (*n* = 3 technical replicates on cDNA obtained using tissue pooled from 2 to 3 plates, each with 20 seedlings **(E)**). Distinct letters indicate significant differences (*P* < 0.05) in two-way ANOVA followed by Tukey’s HSD test (*MYB10*) or two-sample Student’s *t*-tests upon correction for multiple comparisons (*MYB72*). Data in **(F)** are from three independent experiments.

A severe de-regulation in the *irt1* mutant was apparent for a total of 43 Fe deficiency-responsive transcripts of WT, which were not significantly regulated in the same direction in *irt1 v*s. WT (C), and thus showed an unexpected non-responsiveness in *irt1*. These more promising candidate genes for an IRT1-dependent function in root-to-shoot Fe partitioning included *MYB10* and *CYTOCHROME P450 71B5* (*CYP71B5*) (up in WT -Fe) as well as *BETA-GLUCOSIDASE45* (*BGLU45*) and *LACCASE7* (*LAC7*) (down in WT -Fe), for example (Fig. 7A and B, double asterisks; Supplemental Dataset S1). Transcript levels of *MYB10* and *CYP71B5* are known to respond strongly to Fe deficiency (Buckhout et al., 2009; Yang et al., 2010). *BGLU45* and *LAC7* are thought to function in cell wall lignification (Chapelle et al., 2012; Schulten and Krämer, 2017).

We cannot exclude that causative genes may act in shoots, given that *IRT1* expression in phloem companion cells was reported to be critical for its *in planta* function (Marquès-Bueno et al., 2016). In shoots, 76% and 71% of the transcripts that differed in abundance in *irt1* compared to WT under control conditions were also up- or down-regulated, respectively, when WT was exposed to Fe deficiency (Figure 7C, *irt1 vs.* WT (C) ↑ AND - Fe *vs.* C (WT) ↑; Supplemental Dataset S4). Again, some of those transcripts were not up- or down-regulated as expected in Fe-deficient *irt1*, e.g. *ALTERNATIVE OXIDASE1* (*AOX1A*) or *DETOXIFICATION1* (*DTX1*) (Figure 7C and D, double asterisks; Supplemental Dataset S3).

In an independently cultivated experiment, compared to + Fe, the levels of the root-specific transcript *MYB72* were up-regulated *ca.* 100-fold in -Fe-grown WT, but in *irt1* they reached only about 8% of the transcript levels in -Fe-grown WT according to RT-qPCR, thus fully confirming the microarray data (Figure 7E). For *MYB10*, a 10-fold increase in transcript levels was observed in the WT in - Fe, whereas increase in *irt1* was only about 2.6-fold (Fig. 7E, compare with 3-fold change, not statistically significant, according to the microarrays). The *CYP71B5* expression profile was similar to that of *MYB10* and *MYB72* (Supplemental Fig. S8A and E). Thus, according to RT-qPCR, *MYB10* and *CYP71B5* also form part of the regulatory category represented by *MYB72*, so that the microarrays may have been insufficiently sensitive to detect this (Fig. 7A, *). RT-qPCR also suggested that the *bHLH39* expression ratio between *irt1* and -Fe-grown WT was 2.7, whereas the microarray data had suggested a ratio below 2 (Fig. 7A, #; Supplemental Fig. S8B and E). Both microarray and RT-qPCR results supported that root transcript levels of *CYP71B5* were severely under-responsive, and those of *bHLH39* were considerably more responsive, to the general Fe deficiency in *irt1* seedlings. In shoots, fully confirming the microarray data, RT-qPCR detected no statistically significant difference in transcript levels of *NAS4* between WT - Fe and *irt1* + Fe (Supplemental Fig. S8D and E). Transcript levels of *DTX1*, however, in *irt1* were only about 16% of those in WT - Fe (Supplemental Fig. S8C and E), also confirming the microarray data. Thus, in shoots of *irt1*, *DTX1* transcript levels did not follow the physiological Fe status, whereas *NAS4* transcript levels did.

Further consideration of these data highlights some complexities affecting the identification of candidate genes for a role in IRT1-dependent root-to-shoot Fe partitioning. Importantly, the de-regulation of expression of Fe deficiency-responsive genes in the *irt1* mutant could well constitute the cause of altered Fe partitioning in the *irt1* mutant, congruent with candidate gene status. However, the altered expression of such genes in *irt1* could also merely reflect as a consequence the locally altered physiological status of Fe or even of secondary substrate metals in *irt1* by comparison to the WT or of the dynamics of Fe deficiency responses over time. Complicating this even further, the cell-autonomous, non-cell-autonomous or even systemic contributions of the transcriptional regulation of these genes have not been established yet. Finally, the cell type-specificity of candidate gene expression in relation to Fe localization were not considered in our transcriptomics experiment.

To begin to tackle this latter issue, we next attempted to employ information from publicly available cell type-specific gene expression data. Using the GEO2R tool and available cell-type specific datasets (GEO; https://www.ncbi.nlm.nih.gov/geo/, GSE10501), we identified transcripts that are at least 2-fold more abundant either in the cortex or in the endodermis than in the stele under conditions of - Fe (24 h treatment) (Dinneny et al., 2008; Supplemental Figure 9A). Among these genes, we then focused on the subgroup of genes for which our data indicated differential transcript abundance in WT roots between - Fe and + Fe conditions, as putative reporters of local Fe status (Supplemental Dataset S5; Supplemental Dataset S6). However, the expression of many of these genes is likely to be systemically controlled according to the iron status of the shoot, like *IRT1* (Vert et al., 2003). Therefore, among the chosen genes, we finally identified those genes as putative markers of local Fe status, of which transcript levels in *irt1* followed Fe distribution across its roots (See Fig.1B), i.e. did not suggest a more deficient physiological Fe status in *irt1 vs*. WT roots under control conditions. Expression of 38 Fe deficiency-responsive genes is higher in the cortex than in the stele, including *BGLU45*, *ZRT/IRT-LIKE PROTEIN2* (*ZIP2*)*, NAS1, AT3G43670* and *BTSL2*, for example (Dinneny et al., 2008). Additionally, their transcript levels did not show a statistically significant Fe deficiency response in *irt1* compared to WT under control conditions, according to our microarray data from bulk root tissues (Supplemental Dataset S5). We confirmed this by RT-qPCR for *BGLU45* and *BTSL2* (Supplemental Fig. S9B). Given that published data from promoter-GUS fusions of *BTSL2* (Rodríguez-Celma et al., 2019) rather suggest a broad expression domain in Fe-deficient roots, the most promising marker transcript that may reflect local Fe status in root cortex cells remains *BGLU45* encoding beta-glucosidase 45.

### Candidate genes for roles in IRT1-dependent root-to-shoot Fe partitioning

We identified a group of Fe deficiency-responsive genes as under-responsive in roots of the *irt1* mutant (Fig. 7, * and **; Supplemental Datasets S1 and S2). Among these, we identified *MYB72* and *MYB10* as candidates for putative roles in IRT1 protein-dependent root-to-shoot Fe translocation, for example. Therefore, we analyzed transcript levels of *MYB72* and *MYB10* across our panel of transgenic lines, following cultivation as for microarray-based transcriptomics. Indeed, roots of *irt1* S206A lines responded to Fe deficiency, compared to control conditions, with a larger increase of *MYB72* and *MYB10* expression than in *irt1* and *irt1* H232A (Fig. 7F; Supplemental Fig. S10). Additional candidate genes that are also Fe status markers, *NAS4* and *IMA1*, exhibited a similar expression pattern (Supplemental Figure S10). These observations can be interpreted as a partial complementation of an attenuated Fe deficiency responsiveness in *irt1* roots (see Fig. 7). Different from *irt1* S206A, changes in transcript levels between Fe deficiency and Fe replete conditions suggested that the response of *irt1* H232A to Fe levels in the media was overall similarly attenuated as in the *irt1* mutant (Fig. 7). Together, these results are consistent with possible roles of the transcription factor-coding genes *MYB72* and *MYB10* as well as their target gene *NAS4*, and *IMA1*, in IRT1-dependent root-to-shoot Fe partitioning, but further work will be required to obtain unequivocal evidence for such a role of these or other candidate genes.

## DISCUSSION

### High root Fe concentrations in the *irt1* mutant

Our results indicate that roots of the *irt1* mutant accumulate Fe (Fig. 1; Supplemental Fig. S5). By contrast, shoots of the *irt1* mutant – as is well established and confirmed again here – contain reduced Fe concentrations as long as Fe is not growth-limiting, compared to the WT (Figs. 1 and 5). Several earlier studies suggested that the *irt1* mutant may contain unexpectedly high levels of Fe in the root, but the physiological significance of this had remained unaddressed (Henriques et al., 2002; Cailliatte et al., 2010; Ivanov et al., 2014). Indeed, the use of EDTA as a chelator of Fe^3+^ in the synthetic media of these studies favored the precipitation of Fe (Chaney, 1988; Becher et al., 2004; Salomé et al., 2014) and consequently the authors very appropriately considered that an incomplete desorption of apoplastically precipitated or bound Fe could have affected their results. In this study, the use of HBED as Fe^3+^ chelator in combination with a comparably low total Fe concentration are better suited for maintaining Fe^III^ in solution in a synthetic medium. In line with this, Fe concentrations in roots remained in the range of those in shoots of the WT (Fig. 1D).

Our histochemical detection of Fe in *irt1* roots using Perls stain located Fe accumulation predominantly in the cortex of the differentiation zone (Fig. 2). We observed no Fe signal in cell walls in general. Transcript levels of a number of Fe status marker genes indicated an overall more sufficient Fe status in roots than in shoots of the *irt1* mutant, which can be taken to support that the Fe accumulated in roots of *irt1* is at least partly intracellular and physiologically accessible (Figs. 6 and 7, Supplemental Figs. S6 to S10). Our targeted search identified *BGLU45* as a possible marker transcript of local Fe status in root cortex cells that is in agreement with Fe levels in the *irt1* mutant (Supplemental Fig. S9), which will require dedicated confirmation in the future. The physiological Fe status of the shoots systemically governs the transcriptional activation of a number of Fe deficiency-related genes in the roots (Vert et al., 2003; Giehl et al., 2012; Mendoza-Cózatl et al., 2014; Khan et al., 2018). This complicates the identification of cell type-specific markers of local iron status in roots. In the *nramp1 irt1* double mutant Perls-DAB staining localized Fe in the walls of cortex cells, and the authors attributed this observation to the lack of *NRAMP1* function in the *irt1* genetic background (Castaings et al., 2016). The extensive sample preparation carried out in this study may have removed intracellularly localized Fe or deposited it onto cell walls.

When the high-affinity Fe^2+^ transporter IRT1 or the secondary Fe^2+^ transporter NRAMP1 are not present, endodermal suberization is delayed, which was proposed to facilitate the movement of Fe into the stele upon the entry into roots *via* the apoplastic or trans-cellular pathways (Barberon et al., 2016). Despite the decreased suberization and increased permeability of the endodermis for ions in *irt1*, our results support the hypothesis that Fe is immobilized predominantly in the cortex and also in the endodermis of the *irt1* mutant, i.e., partly outward from the endodermis. This is consistent with our hypothesis that the inward transport of Fe^2+^ across the plasma membrane of root epidermal cells is not the only function of the IRT1 protein.

### Phenotypic rescue of the root-to-shoot Fe partitioning defect of *irt1* by the transport-inactive mutant IRT1_S206A_ variant

Based on the results presented here, we hypothesized that the IRT1 protein acts indirectly to de-repress root-to-shoot partitioning of Fe, in addition to its well-known direct membrane transporter function of mediating the uptake of Fe^2+^ into root cells. If our hypothesis is correct, we would expect to observe the partial complementation of *irt1* mutant phenotypes by an IRT1 variant that is incapable of transporting Fe^2+^ but maintains indirect functionality. While the latter is entirely unknown to date, the direct transmembrane ion transport function of IRT1 is very well-studied. Amino acid residues Serine 206 and Histidine 232 are both critical for the cellular import of Fe^2+^ (Fig. 4, Supplemental Fig. S2), as well as of secondary substrates Zn^2+^, Mn^2+^ and likely also Cd^2+^, conferred by IRT1 in the heterologous system *Saccharomyces cerevisiae* (Eide et al., 1996; Rogers et al., 2000).

We introduced wild-type IRT1, IRT1_S206A_ and IRT1_H232A_ into an *irt1* mutant background and then tested for the rescue of various aspects of phenotype of Arabidopsis *irt1* mutants. Our results are in agreement with our hypothesis, and they indicated that IRT1_S206A_ is inactive with respect to the direct function of IRT1, but maintains the indirect function at least partially. By contrast, our results suggest that IRT1_H232A_ is produced and stable *in planta*, but maintains neither the direct nor the indirect function of IRT1. We came to these conclusions by quantifying Fe concentrations and Fe deficiency symptoms in roots and shoots of seedlings (Figs. 3 - 7, Supplemental Figs. S3 to S8). Important support came from the quantification of Mn concentrations in tissues, exploiting the known transport activity of IRT1 for the secondary substrate Mn^2+^ in combination with IRT1 protein abundance under deficient and replete physiological Fe status (Figs. 3 and 5, Supplemental Figs. S3 and S5; Vert et al., 2002).

Intriguingly, we were unable to identify any difference between WT and *irt1* roots in short-term Fe accumulation, i.e. after 15 min of exposure to 2 μM radiolabeled Fe^2+^ (Fig. 4B, Supplemental Table S2). Previous evidence from short-term Fe uptake assays *in planta* is scarce. In contrast to 2-w-old Fe-deficient WT plants, the *irt1* mutant (Ws background) was incapable of accumulating ^55^Fe in shoots after 48 hours (Vert et al., 2002). Wild-type roots contained approximately 50% and *irt1*-3 roots 70% of the total Fe accumulated per plant upon 1 h of pulse-labeling with 3 μM ^59^Fe-labeled Fe^2+^ *via* the roots of 2-w-old seedlings cultivated in Fe- and Zn-deficient medium. There was large quantitative variation and total Fe uptake averaged of 130 ± 60 and 80 ± 30 fmol Fe plant^−1^ in WT and *irt1*-3, respectively, and images suggested an overall smaller size of *irt1*-3 than of WT seedlings used in the pulse labeling experiments (Henriques et al., 2002; T-DNA insertion 133 bp upstream of translational start codon). Although a direct or quantitative comparison with these two earlier studies is impossible, our results are in general agreement. Taking these studies together, the unequivocal defect in *irt1* mutants is a dramatically reduced Fe accumulation rate in shoots, whereas root Fe accumulation rate in *irt1* is unchanged or even slightly enhanced. Thus, *irt1* mutants retain some root Fe uptake capacity independent of the IRT1 protein, suggesting that secondary Fe uptake systems operate in these mutants under the experimental conditions employed. An involvement of NRAMP1, for example, is possible, as proposed (Cailliatte et al., 2010; Castaings et al., 2016).

### Candidate genes for a role in IRT1-dependent Fe partitioning from the root to the shoot

Our results support the hypothesis of a secondary function of IRT1 in the partitioning of Fe from the root to the shoot, which is at least partially independent of the primary function of IRT1 in root cellular Fe uptake. Such a secondary function of IRT1 could involve a physical interaction of IRT1 with an unknown protein, which then triggers a process that finally results in the radial passage of Fe from root cortex cells inwards, for example. According to a symplastic pathway model, this IRT1-dependent process could encompass maintaining symplastic mobility of Fe through its chelation in the cytosol, suppressing the immobilization of Fe in vacuoles of root cortex cells, or fostering symplastic radial cell-to-cell passage of Fe *via* plasmodesmata. In a transcellular pathway model, the envisaged process could involve cellular Fe efflux from root cortex cells and Fe re-uptake by endodermal cells. Proteins mediating any of these processes in a localized fashion remain unidentified. IRT1-dependent de-repression of root-to-shoot partitioning may occur entirely through translational or post-translational regulation, or it may involve changes in transcript levels of some of the involved genes. Here we searched for candidate Fe deficiency-responsive transcripts that were under-responsive or non-responsive in roots of the *irt1* mutant.

By comparison to Fe-deficient WT, in roots of *irt1* we observed lower expression of several FIT-dependent (e.g. *CYP71B5, BGLU42*) and FIT-independent (e.g. *NAS4, bHLH101*) genes, among others (Supplemental Datasets S1 and S3; Colangelo and Guerinot, 2004; Bauer et al., 2007; Zamioudis et al., 2015; Schwarz and Bauer, 2020). Notably, the Fe deficiency-responsive increase in *MYB10* and *MYB72* transcript levels was strongly attenuated in *irt1* (Fig. 7). Root-specific transcription factors MYB72 and MYB10 regulate Fe partitioning to the shoot *via* activating *NAS4* transcription and thus NA biosynthesis in the stele (Palmer et al., 2013). Later, MYB72 was found to also activate the transcription of genes encoding the enzymes F6’H1 and BGLU42 of the coumarin biosynthesis pathway that contribute to Fe mobilization in soil (Schmid et al., 2014; Zamioudis et al., 2014; Stringlis et al., 2018). In summary, candidate genes *MYB72* and *MYB10*, and indirectly *NAS4*, may well account for impaired root-to-shoot partitioning of Fe in *irt1*. However, for these and a number of other potential candidate genes (Supplemental Datasets S1 to S4), we were unable to fully resolve whether their de-regulation is the cause or the effect of Fe immobilization in *irt1* roots.

## CONCLUSIONS

Here we provide evidence for a secondary function of IRT1 in the partitioning of Fe from the root to the shoot. The ability of a transport-inactive IRT1_S206A_ variant to partially complement the *irt1* mutant supports our hypothesis and indicates that the secondary function of IRT1 is at least partly independent of its primary function of high-affinity Fe uptake into root epidermal cells. We identify previously characterized genes as candidate downstream targets of IRT1-dependent transcriptional regulation, which could contribute to Fe mobilization towards the shoot. Future work will be directed at the signaling and mechanisms of IRT1-dependent Fe partitioning from roots to shoots.

## MATERIALS AND METHODS

### Plant material and growth conditions

All plant lines used in the present study are in the Col-0 background. WT seeds were obtained from NASC. The *irt1*-2 mutant (*pam42*) was from Varotto et al. (2002). The transgenic lines *irt1 IRT1_P_:IRT1* (WT *IRT1*), *irt1 IRT1_P_:IRT1S206A* (*irt1* S206A) and *irt1 IRT1_P_:IRT1H232A* (*irt1* H232A) were generated herein.

For sterile growth, seeds were washed in 70% (v/v) ethanol for 1 min, surface-sterilized with 1.4% (w/v) NaOCl and 0.02% (v/v) Triton X-100 for 10 min and washed five times with ultrapure water (Milli-Q; Merck). Following seed stratification at 4°C for 2 d, seedlings were pre-cultivated on a modified Hoagland’s solution containing macro- and micronutrients as used in Haydon et al. (2012), supplemented with 1% (w/v) sucrose and solidified with 0.8% (w/v) agar Type M (Sigma). Ten-d-old seedlings were then transferred using titanium tweezers to modified Hoagland’s solution supplemented with 1% (w/v) sucrose in which Fe was omitted from the nutrient solutions (− Fe treatment) or supplied as 10 μM FeHBED (+ Fe treatment). Treatment media were solidified with 0.8% (w/v) agar Type M (Sigma) depleted from contaminant metals by EDTA-washing as described in Haydon et al. (2012). Seedlings were grown on vertically oriented square 120-mm polystyrene Petri dishes (Greiner Bio-One) in 11-h light (22°C, 145 μmol m^−2^ s^−1^ white light) and 13-h dark (18°C) cycles (Percival CU-41L4; CLF Climatics). For selection of homozygous transgenic lines carrying a single T-DNA insertion, seeds were grown on 0.5x Murashige & Skoog (MS) medium (Duchefa) supplemented with 1% (w/v) sucrose, 0.8% (w/v) agar and hygromycin (30 μg mL^−1^). In the T2 generation, five to six independent lines per construct were grown on MM media (Haydon and Cobbett, 2007) containing 1% (w/v) sucrose and 0.8% (w/v) agar Type M supplemented with hygromycin (30 μg mL^−1^) for 10 d and then transferred to treatment media (+ Fe treatment, as described above) for an additional 17 d before harvest (see Supplemental Table S1). For hydroponic plant cultivation for short-term uptake assays of radiolabeled Fe, seeds were surface-sterilized, sown on 0.5x modified Hoagland’s solution supplemented with 5 μM FeHBED (WT and *irt1* IRT1) or 10 μM FeHBED (*irt1*, *irt1* S206A and *irt1* H232A) and solidified with 0.65% (w/v) Noble agar (Sigma) in 0.7 mL black Eppendorf tubes and stratified at 4°C for 4 d. At the age of 7 d, the bottom third of each tube was cut off and about 60 tubes per line (1 seedling per tube) were transferred to a styrofoam tray floating on 4 L of 0.5x modified Hoagland’s solution supplemented with 10 μM FeHBED (WT and *irt1* IRT1) or 50 μM FeHBED (*irt1*, *irt1* S206A and *irt1* H232A) in a box covered with Saran wrap. After 1 week, plants were carefully transferred to smaller boxes (4 plants per box in 420 mL solution, keeping roots separate per plant) containing modified Hoagland’s solution supplemented with FeHBED as mentioned above (pre-cultivation). Plants were grown for an additional 5 weeks (WT and *irt1* IRT1) to 6 weeks (*irt1*, *irt1* S206A and *irt1* H232A) with a weekly exchange of solutions to obtain vegetative stage plants of similar sizes. Roots were subsequently washed in 10 μM EDTA for 2 min and transferred into modified Hoagland’s solution lacking FeHBED for 6 days (Fe deficiency treatment). Plants were grown in 11-h light (20°C, 145 μmol m^−2^ s^−1^ white light) and 13-h dark (18°C) cycles (Percival CU-41L4; CLF Climatics) in a tray that was covered with a transparent cover, which was removed during the final 3 weeks of cultivation. For the confirmation experiment at MU, pre-cultivation was in 5 μM (WT) and 50 μM FeHBED (*irt1*) for 4 to 5 w, and Fe deficiency treatment was in 0 μM (WT) and 5 μM (*irt1*) FeHBED for 5 d. For soil cultivation, seeds were germinated on Minitray soil (Balster Einheitserdewerk, Fröndenberg) after 4 days of stratification at 4°C. Plants were grown under the same environmental conditions as for sterile growth for 3 weeks.

### Yeast Constructs, Strains, and Growth

The *IRT1* coding cDNA sequence, including translational start and stop codon, was amplified from the corresponding transgenic plant line by RT-PCR with IRT1-ATG_F 5’-CACCATGGCTTCAAATTCAGCAC-3’ and IRT1-cDNA_TAA_R 5’- TTAAGCCCATTTGGCGATAATCG-3’, introduced into pENTR-D/TOPO (Invitrogen) and cloned into the yeast expression vector pFL61-Gateway (Desbrosses-Fonrouge et al., 2005) using LR clonase (Invitrogen) according to the manufacturer’s protocol. Mutations S206A and H232A were introduced by a modified QuikChange protocol (Zheng et al., 2004) using KAPAHiFi DNA polymerase (PEQLAB Biotechnology) and the primers AtIRT1_S198A_For 5’- TAGTTCACGCGGTGGTCATTGGATTATC-3’, AtIRT1_S198A_Rev 5’- CCACCGCGTGAACTATGATCCCAAG-3’, AtIRT1_H224A_For 5’- TGCTTCGCTCAAATGTTCGAAGGCATGGG-3’ and AtIRT1_H224A_Rev 5’- CATTTGAGCGAAGCAAAGAGCTGCTATAAGTC3’. All constructs were verified by Sanger sequencing. For complementation assays, competent *S. cerevisiae* yeast cells from WT (DY1457) and *fet3fet4* mutant (DEY1453; Dix et al., 1994) strains were transformed with each construct or the empty vector (Dohmen et al., 1991). As a positive control, the corresponding WT yeast strain was transformed with the empty vector. Transformants were selected on SD media lacking uracil (SD-Ura) pH 5.7, 2% (w/v) Bacto-Agar, with 2% (w/v) D-glucose as a carbon source. For each construct, three independent transformant colonies were grown overnight at 30°C in 2 mL liquid SD-Ura pH 3.5 with 2% (w/v) D-glucose supplemented with 10 μM FeCl_3_, to early stationary phase (OD_600_ ≈ 0.3, *ca*. 10^7^ cells ml^−1^). Yeast cells were then centrifuged at 15,700x*g* for 1 min, washed once in SD-Ura pH 5.7 with 2% (w/v) D-glucose to remove residual Fe and subsequently resuspended in the same medium. Aliquots of 10 μL of ten-fold serially diluted cell suspensions (OD_600_ of 0.3, 0.03,…) were spotted onto SD-Ura pH 5.7 containing 2% (w/v) Bacto-Agar and 2% (w/v) D-glucose, supplemented with 0.5 mM FeSO_4_ or left unamended. Plates were incubated at 30°C for 3 days and photographed using a Nikon Digital SLR camera with an AF-S DX Zoom NIKKOR 18-70 mm 1:3,5-4,5G ED-IF objective.

### Generation of transgenic plants

For genetic complementation of *irt1* mutant with wild-type *IRT1*, the promoter region (1,024 bp upstream of the translational start codon), the coding region, and the 3’ UTR sequence (310 bp downstream of the stop codon) were amplified by PCR using Phusion DNA polymerase (NEB) from genomic Col-0 DNA with IRT1p_-1024F 5’- CACCGACACATTAAACATTCATACCCGATT-3’ and IRT1_1546R 5’- CTTTAATTTACTTATCTTGAAAAAGCAGC-3’. For the mutant variants *IRT1_S206A_* and *IRT1_H232A_*, overlapping PCR products were generated with IRT1p_-1024F and IRT1_S198A_R 5’- GATAATCCAATGACCACCGCGTGAACTATGATCCCAAG-3’ or IRT1_H224A_R 5’- CCATGCCTTCGAACATTTGAGCGAAGCAAAGAGCTGC-3’ and IRT1_1546R and IRT1_S198A_F 5’-CTTGGGATCATAGTTCACGCGGTGGTCATTGGATTATC-3’ or IRT1_H224A_F 5’ GCAGCTCTTTGCTTCGCTCAAATGTTCGAAGGCATGG-3’. These PCR products were combined in a subsequent PCR reaction using only IRT1_-1024F and IRT_1546R to generate the full-length mutant variants. WT and mutant PCR products were introduced into pENTR-D/TOPO (Invitrogen) and verified by Sanger sequencing before recombining into a modified pMDC32 vector (Curtis and Grossniklaus, 2003) lacking the 35S promoter using LR Clonase (Invitrogen) (Hanikenne et al., 2008). Plasmids were introduced into *Agrobacterium tumefaciens* GV3101 by electroporation and transformed into *irt1* by floral dip (Clough and Bent, 1998). Homozygous T3 seeds carrying single-locus insertions of two independent transgenic lines per construct were chosen for detailed characterization based on shoot Fe concentrations in the T2 generation (Supplemental Table S1).

### Microarray-based transcriptomics

All methods in this Section are described according MIAME recommendations. Fifteen-d-old seedlings were used for the microarray experiment. Roots and shoots were harvested separately at ZT 3 h, flash-frozen in liquid nitrogen and subsequently stored at −80°C. RNA was extracted using RNeasy Plant Mini Kit with on-column DNase I digestion (Qiagen). cRNA labeling and ATH1 hybridization were performed according to manufacturer’s instructions at GeneCore Facility, EMBL, Heidelberg. The array GeneChip ATH1-121501 (Thermo Fisher Scientific, former Affymetrix), which contains 22,500 probe sets representing approximately 24,000 gene sequences of the *A. thaliana* genome, was used for hybridizations. Replicate ATH1 arrays were hybridized with labeled cRNA from two independent experiments, with six samples per experiment: shoots or roots of WT + Fe, WT -Fe and *irt1* + Fe. Signal intensities were imported into Genespring (Version 8; Agilent Technologies), and an RMA normalization was performed without transformation. Differentially expressed genes (DEG) were selected by ≥ 2-fold difference in mean normalized signal in both replicates and an unpaired Mann-Whitney test with Benjamini-Hochberg adjustments for multiple comparisons, with a cutoff value of *P* < 0.15. The gene lists obtained with different comparisons were then intersected with one another to identify overlapping DEGs using Venn Selector at VIB (UGent; http://bioinformatics.psb.ugent.be/webtools/Venn).

### RT-qPCR

For gene expression analysis using RT-qPCR, seedlings were grown and harvested as for the microarray experiments. Total RNA was extracted from 30 to 50 mg of frozen powdered tissue with Trizol Reagent following the manufacturers’ instructions. Total RNA was treated with 2 U of DNase per 10 μg of total RNA using the TURBO DNA-free™ Kit (Thermo Fisher Scientific, former Ambion). Integrity of DNA-free RNA was verified in Bleach Gel (Aranda et al., 2012), and A280/A260 ratio in NanoDrop™ 2000/2000c (Thermo Fisher Scientific) was used to assess RNA quality. One μg of DNase-treated RNA and oligo (dT)_18_ primers were used in first-strand cDNA synthesis with the RevertAid First Strand cDNA Synthesis Kit (Thermo Fisher Scientific) following manufacturer’ instructions. RT-qPCR was performed on a LightCycler480 (Roche) in a 10-μL reaction mixture containing 8 ng of cDNA, each primer at 0.25 μM and 5 μL of 2X GoTaq qPCR Master Mix (Promega). The amplification program consisted of preincubation steps at 50°C for 2 min and 95°C for 10 min followed by 40 cycles of 95°C for 15 s and 60°C for 1 min. A final dissociation step of 95°C for 10 s and 65°C for 5 s was performed for melting curve analysis. Reaction efficiencies (E) for each PCR reaction were determined with the LinRegPCR program, version 2016.0 (Ruijter et al., 2009) and used to calculate Transcript Level as TL = E^−Ct^, with Ct as the cycle threshold. Relative TL (RTL) was calculated by dividing TL of the gene of interest by geometric mean of TLs of two housekeeping genes, *UBQ10* and *EF1α*. All oligonucleotide sequences are specified in Supplemental Table S3.

### Ionomics and biomass estimates

Twenty-d-old seedlings were used for the quantification of metal concentrations. Shoots and roots were harvested separately by cutting below the hypocotyl with a scalpel washed in 10 mM EDTA. Root and shoot tissues were pooled per plate and desorbed by washing in 5 mM CaSO_4_, 10 mM MES-KOH, pH 5.7 for 10 min, in 5 mM CaSO_4_, 10 mM Na_2_EDTA, pH 5.7 for 5 min, and twice in ultrapure water for 1 min, all carried out on ice with occasional stirring. After desorbing, tissue pools were blotted dry using paper towels (Blauer Engel), and fresh biomass (FW) was recorded and calculated per plant. Aliquots of 4 to 25 mg of powdered dry biomass per sample were digested as described in Sinclair et al. (2017). Multi-element analysis was conducted using Inductively Coupled Plasma Optical Emission Spectrometry (ICP-OES) in an iCAPDuo 6500 instrument (Thermo Fisher Scientific), following calibration with a blank and a series of five multi-element standards from single-element standard solutions (AAS Standards; Bernd Kraft). The precision of measurements was validated by measuring a sample blank and an intermediate calibration standard solution, as well as digests of a certified reference material (Virginia tobacco leaves, INCT-PVTL 6; Institute of Nuclear Chemistry and Technology, PL) before and after every sample series. To ensure timely handling of the samples, we divided the panel of transgenic lines into two sets, each representing all constructs, and we used in each experiment one set plus WT and *irt1*.

### Whole-mount staining of non-heme Fe^III^ with Perls reagent

Seven-d-old seedlings germinated and grown in standard media were harvested at ZT 1 h. Three seedlings were pooled in a 2-mL tube, washed once with ice-cold 10 mM EDTA pH 5.7 for 5 min and three times with ice-cold ultrapure water for 1 to 2 min. Seedlings were then submerged in 1 mL Perls solution (2% (v/v) HCl and 2% (w/v) potassium ferrocyanide), vacuum infiltrated for 15 min (500 mbar) and incubated at room temperature for 30 min (Roschzttardtz et al., 2009). Seedlings were then rinsed twice with ultrapure water and mounted in ultrapure water for visualization on an Imager.M2 (Zeiss).

### Chlorophyll and root length measurements

Quantification of leaf chlorophyll content and root length were done on 15-d-old seedlings. For chlorophyll measurements, 34 to 52 mg of pooled shoots (5 to 8 seedlings) were harvested into a 2-mL tube, flash-frozen in liquid nitrogen and homogenized with a plastic pistil cooled in liquid nitrogen. After the addition of 1.5 mL 100% (v/v) methanol (Sigma), samples were covered with aluminum foil and incubated at 70°C for 15 min, stirring at 850 rpm for 10 sec every 5 min. After cooling on ice for 5 min, methanol extracts were cleared by centrifugation at 16,000x*g*, 4°C, for 1 min. Absorbance (extinction) values were measured at 652 nm and 665 nm in a Synergy HTX Multi-Mode Reader (Agilent, former BioTek), using methanol as a blank. Microplate pathlength correction was applied according Warren (2008). Chlorophyll concentrations were calculated following Porra et al. (1989). Three repeated measurements of each sample served as technical replicates. For root elongation estimates, the positions of primary root tips of 10-d-old seedlings were marked with a permanent marker (Edding® 400) on the bottom of the petri plate immediately after transfer to + Fe or - Fe treatment media. Five days afterwards, seedlings were imaged with a Nikon Digtal SLR camera with an AF-S DX Zoom NIKKOR 18-70 mm 1:3,5-4,5G ED-IF objective at 50 mm distance. Primary root length was measured using ImageJ (Schneider et al., 2012) by drawing a segmented line along the main root.

### Root surface Ferric Chelate Reductase (FCR) activity assays

Seedlings were grown as for microarray experiments. Roots of 5 to 8 15-d-old seedlings (8 to 33 mg fresh biomass) were harvested at ZT 3 h using a scalpel, pooled and immediately transferred to 2-mL tubes securely capped to avoid humidity loss. Roots were incubated in 1.5 mL of assay solution (0.1 mM Fe^III^EDTA, 0.3 mM FerroZine) at RT in darkness for 20 min (time course experiment) or 40 min (FCR activity after Fe deficiency for 5 d). After reading absorbance at 535 nm on a PowerWave XS2 plate reader (Agilent, former BioTek), Fe^II^FerroZine concentrations were calculated using an extinction coefficient of 28.6 mM^−1^ cm^−1^ (Gibbs, 1976).

### IRT1 Immunoblots

Total protein was extracted from 30 mg FW of pooled root tissues as described earlier (Sgula et al., 2008) with the difference that 2% (w/v) DTT was used as reducing agent and that prior to centrifugation, protein extracts were incubated 5 min at room temperature (RT) to facilitate membrane solubilization. Total protein in the supernatants was quantified in 1:5 dilutions with the Pierce™ BCA Protein Assay Kit using BSA as a standard. Per sample, 20 μg of total protein was mixed with ultrapure water and 2 μL of 6x loading buffer (375 mM Tris-HCl (pH 6.8), 30% (v/v) mercaptoethanol, 0.03% (v/v) bromophenol blue, 12% (w/v) SDS, and 60% (v/v) glycerol) in a final volume of 12 μL and subsequently incubated for 20 min at 37°C. Proteins were separated by 12% (w/v) SDS-PAGE and blotted onto a polyvinylidene difluoride (PVDF) membrane by wet/tank transfer (Towbin et al., 1979). The PVDF membrane was blocked with 5% (w/v) skim milk in TBS-T (0.05% Tween-20) at RT for 1 h and probed with anti-IRT1 antibody (1:5,000, Agrisera AS111780, Lot number 1203) in 2.5% (w/v) skim milk in TBS-T (0.05% Tween-20) at RT for 1 h. Following incubation at RT for 1 h with the secondary antibody goat anti-rabbit IgG (H+L), HRP (1:20,000, Thermo Fisher Scientific 31466; lot number RL243150) in 2.5% (w/v) skim milk in TBS-T, signals were detected with ECL Select Western Blotting Detection Reagent (GE Healthcare) and a Fusion Fx7 documentation system (Vilber Lourmat).

### Short-term uptake rates of ^55^Fe

Hydroponically cultivated 7.5- (WT and *irt1* IRT1) or 8.5- (*irt1*, *irt1* S206A and *irt1* H232A) week-old plants of equal sizes were used for short-term Fe uptake assays. To allow for accurate sample processing we included one line per construct, WT and *irt1* in each experiment. At ZT 1 h, the root systems of 4 to 5 replicate plants per line were submerged in glass liquid scintillation vials (one plant per vial) containing 35 mL of 1 mM MES-KOH pH 5.7, 2.1 μM citrate, 1 mM ascorbic acid and 2 μM ^55^Fe (^55^FeCl_3,_ 0.0197 MBq nmol^−1^, Perkin-Elmer). After two incubations of 1.5 min and 15 min duration for each genotype, respectively, roots were desorbed by washing 3 times in fresh ice-cold solution of 5 mM CaSO_4_, 10 mM Na_2_EDTA (pH 5.7), 1 mM MES-KOH (pH 5.7) for 5 min (40 mL per plant). Roots were cut off, carefully blotted dry between paper towels (Blauer Engel), weighed and stored at −20°C for 24 hours in 6 ml HPDE scintillation vials to destroy the cells. Roots were then suspended in 4 mL of a scintillation cocktail at RT (Rotiszint, Carl Roth, Germany). Counting was performed twice for each sample (10 min per sample) with a ß-scintillation counter Beckman LS6000TA (Beckman Coulter, USA). Roots of two plants identically grown and processed but not exposed to the incubation solution were used to determine background counts. For calculation of ^55^Fe uptake, values of uptake after 1.5 min were subtracted from those after 15 min uptake and expressed as nmol ^55^Fe g^−1^ FW h^−1^. Pre-cultivation periods and conditions were slightly varied from experiment to experiment, gradually obtaining plants of more and more equivalent sizes and developmental stages for all genotypes. For a confirmation experiment at MU (Supplemental Table S2), plants of similar size grown hydroponically were pre-incubated in 35 mL of a solution of 0.2 mM CaSO_4_, 1 mM ascorbic acid, 5 mM MES-TRIS, pH 5.0, for 30 min, and then transferred to 30 mL of uptake buffer of identical composition with the addition of 1 μM FeCl_3_ (spiked with *ca.* 2 μCi ^59^Fe) for an uptake period of 120 min. Washing solutions were composed as follows: 20 mM TRIS, 5 mM EDTA (30 mL for 10 min, followed by 35 mL for 10 min), ultrapure water (25 mL for 5 min, twice). Roots were separated from shoots and blotted dry as described above before scintillation counting. Vials containing only scintillation cocktail were used for background subtraction.

### Statistical analysis

Statistical analyses were conducted in R Studio Version 1.1.456 (R Core Team, 2020). Two-factor ANOVA was performed assuming the following linear model: model = lm (Var1 ~ TREATMENT + GENOTYPE + TREATMENT:GENOTYPE). Tukey HSD was applied in multiple comparisons of means. Alternatively, when homoscedasticity was not met (*P* < 0.05, Levene’s test), pairwise Welch *t*-tests were conducted. When normality was not met (*P* < 0.05, Shapiro-Wilk test), pairwise Student’s *t*-tests were used. For all *t*-tests, pairwise differences were considered significant when FDR-corrected *q* < 0.05.

### Accession numbers

Sequence data for the genes mentioned in the text can be found on The Arabidopsis Genome Initiative (TAIR) or GenBank website. Accession numbers are (in alphabetic order): *ALTERNATIVE OXIDASE1A* (*AOX1A*), *AT3G22370*; *basic HELIX LOOP HELIX39* (*bHLH39*), *AT3G56980*; *basic HELIX LOOP HELIX 101* (*bHLH101*), *AT5G04150*; *BETA-GLUCOSIDASE42* (*BGLU42*), *AT5G36890*; *BETA-GLUCOSIDASE45* (*BGLU45*), *BRUTUS-LIKE2* (*BTSL2*), *AT1G18910*; *AT1G61810*; *CYTOCHROME P45071b5* (*CYP71B5*), *AT3G53280*; *DETOXIFICATION1* (*DTX1*), *AT2G04040*; *FERRITIN1* (*FER1*), *AT5G01600*; *IRONMAN1* (*IMA1*), *AT1G47400*; *IRON REGULATED TRANSPORTER1* (*IRT1*), *AT4G19690*; *IRON REGULATED TRANSPORTER2* (*IRT2*), *AT4G19680*; *MYB DOMAIN PROTEIN10* (*MYB10*), *AT3G12820*; *MYB DOMAIN PROTEIN 72* (*MYB72*), *AT1G56160*; *NICOTIANAMINE SYNTHASE4* (*NAS4*), *AT1G56430*; *ZRT/IRT-LIKE PROTEIN 2* (*ZIP2*), *AT5G59520*; *ZINC-INDUCED FACILITATOR1* (*ZIF1*), *AT5G13740*. The microarray data set accession number is xxx (https://www.ncbi.nlm.nih.gov/geo/).

## SUPPLEMENTAL MATERIAL

**Supplemental Figure S1.** Details of the *irt1* phenotype.

**Supplemental Figure S2.** Topology model of IRT1 highlighting the amino acid residues mutated in this paper.

**Supplemental Figure S3.** Details of *IRT1* expression in transport-inactive mutants.

**Supplemental Figure S4.** Photographs of Arabidopsis WT, *irt1*, *irt1* IRT1, *irt1* S206A, *irt1* H232A plants.

**Supplemental Figure S5.** Root Fe concentration, total Fe per plant, root Mn concentration and *NRAMP1* expression.

**Supplemental Figure S6.** Fe deficiency responses are not constitutively activated in shoots of *irt1* S206A lines.

**Supplemental Figure S7.** Experimental rationale of transcriptomic analysis.

**Supplemental Figure S8.** Validation of microarray data using RT-qPCR.

**Supplemental Figure S9.** Lack of Fe-deficiency response in cortex-enriched transcripts in the *irt1* mutant.

**Supplemental Figure S10.** Transcript levels of candidate genes for roles in IRT1-dependent root-to-shoot translocation across genotypes.

**Supplemental Table S1.** Fe concentrations in shoots of independent transgenic lines for *irt1* IRT1, *irt1* S206A and *irt1* H232A.

**Supplemental Table S2.** Short-term uptake of radiolabeled Fe into roots of Fe-deficient wild-type and *irt1* mutant plants.

**Supplemental Table S3.** Oligonucleotides used in the present study.

## LARGE DATASETS

**Supplemental Dataset S1.** List of overlapping differentially expressed genes between -Fe *vs*. C (WT) and *irt1* (C) *vs*. WT (-Fe) in roots (**, *; Fig. 7A, B).

**Supplemental Dataset S2.** List of differentially expressed genes that overlap between -Fe *vs*. C (WT) and *irt1 vs*. WT (C) in roots (#; Fig. 7A, B).

**Supplemental Dataset S3.** List of overlapping differentially expressed genes between -Fe *vs*. C (WT) and *irt1* (C) *vs*. WT (-Fe) in shoots (**, *; Fig. 7C, D).

**Supplemental Dataset S4.** List of differentially expressed genes that overlap between -Fe *vs*. C (WT) and *irt1 vs*. WT (C) in shoots (#; Fig. 7C, D).

**Supplemental Dataset S5.** List of transcripts reported at higher or lower abundance in the stele *vs*. the cortex that were Fe deficiency-responsive according to this study.

**Supplemental Dataset S6.** List of transcripts reported at higher or lower abundance in the stele *vs*. the endodermis that were Fe deficiency-responsive according to this study.

## Acknowledgments

The *fet3fet4* and corresponding wild-type yeast strains were kindly provided by Prof. Dr. David Eide (University of Wisconsin-Madison, US), Arabidopsis *pam42* by Prof. Dr. Dario Leister (LM University Munich, Germany). We are grateful to Petra Düchting for multi-element analysis, Dr. Lara Syllwasschy for t-test scripts, Andreas Aufermann and Iris Sandorf for technical assistance in plant cultivation, and all lab members for comments (Ruhr University Bochum, Germany). We also thank María de los Ángeles and Patricia Lorente for technical support, and Rafael Picorel for providing infrastructure (EEAD, CSIC, Spain).

