## Supplemental Tables and Figures for "Root-to-shoot iron partitioning in Arabidopsis requires IRON-REGULATED TRANSPORTER1 (IRT1)"

**Supplemental Table S1. Fe concentrations in shoots of independent transgenic lines for *irt1* IRT1, *irt1* S206A and *irt1* H232A.** Data for the transgenic lines are from hygromycin-resistant seedlings in the T2 generation.

| **Genotype** | **Line no.** | **Fe concentration in shoot ($\mu g\ g^{-1}$ DW)** | **Significant difference from *irt1*** | **Significant difference from WT** |
|---|---|---|---|---|
| | | mean ± SD ($n = 3^{§}$) | | |
| WT | | 80.33 ± 5.44 | ** | |
| *irt1* | | 48.32 ± 5.73 | | ** |
| *irt1* IRT1 | i (-1)$ | 102 ± 10.7 | ** | |
| | iii | 103 ± 22.8 | * | |
| | iv | 116 ± 12.2 | ** | * |
| | v (-2)$ | 108 ± 4.8 | *** | * |
| | vi | 105 ± 1.3 | *** | ** |
| | vii | 63 ± 2.9 | * | * |
| *irt1* $IRT1_{S206A}$ | ii (-1)$ | 77 ± 3.2 | * | |
| | iii | 78 ± 6.3 | * | |
| | iv | 138 ± 40.1 | * | |
| | vi | 100 ± 6.5 | ** | |
| | vii (-2)$ | 84 ± 12.3 | * | |
| *irt1* $IRT1_{H232A}$ | i | 46 ± 4.2 | | * |
| | ii (-1)$ | 58 ± 1.5 | | * |
| | iii | 49 ± 8.3 | | * |
| | iv (-2)$ | 50 ± 5.5 | | * |
| | vi | 70 ± 8.1 | | |
| | vii | 52 ± 1.9 | | * |

*$P < 0.05$, **$P < 0.01$, ***$P < 0.001$ (two-sample *t*-tests with corrections for multiple comparisons).

$ Lines characterized in detail in this paper.

§ One replicate comprised pooled shoots of *ca*. 20 seedlings from one petri plate, cultivated under Fe-sufficient conditions (see Materials and Methods).

**Supplemental Table S2.** Short-term uptake of radiolabeled Fe into roots of Fe-deficient wild-type and *irt1* mutant plants.

| Genotype | | **Fe concentration** (nmol Fe $g^{-1}$ FW) | | | | |
|---|---|---|---|---|---|---|
| | | **mean ± SD** | | | | |
| | | **Exp. 1 - Krämer Lab** | | **Exp. 2 - Krämer Lab** | | **Exp. 3 - Mendoza Lab** |
| | *Uptake period* | *5 min* | *15 min* | *1.5 min* | *15 min* | *120 min* |
| WT | | 7 ± 1.8 | 33 ± 24.1 | 3.2 ± 3.2 | 48 ± 49.4 | 31 ± 4.6 |
| *irt1* | | 16 ± 4.9 | 22 ± 8.4 | 7.4 ± 9.4 | 43.6 ± 14.0 | 42 ± 9.4 |

**Exp. 1 and 2**: 2 µM Fe in uptake solution, $n$ = 4 to 5 plants (Fig. 4B is based on Exp. 2); **Exp. 3**: 1 µM Fe in uptake solution, $n$ = 3 plants, shoot: 0 to 0.3 nmol Fe $g^{-1}$ FW (fresh biomass).

**Supplemental Table S3.** Oligonucleotides used in the present study.

| Oligo name | Gene | Sequence (5'- 3´) | Reference |
|---|---|---|---|
| EF1a_qRT_f<br>EF1a_qRT_r | *EF1α* | TGAGCACGCTCTTCTTGCTTTCA<br>GGTGGTGGCATCCATCTTGTTACA | Czechowski et al., 2005 |
| qRT_UBQ10_Fw<br>qRT_UBQ10_Rv | *UBQ10* | GGCCTTGTATAATCCCTGATGAATAAG<br>AAAGAGATAACAGGAACGGAAACATAGT | Haydon et al., 2012 |
| qRT_IRT1_Fw<br>qRT_IRT1_Rv | *IRT1* | CCCCGCAAATGATGTTACCTT<br>GGTATCGCAAGAGCTGTGCAT | This work |
| qRT_IRT2_Fw<br>qRT_IRT2_Rv | *IRT2* | CGTAGCCATTGTTGCCATA<br>GGTTCCAAGGATGATTCCAG | Vert et al., 2009 |
| qRT_FER1_Fw<br>qRT_FER1_Rv | *FER1* | TCCCCAGTTAGCTGATTTCG<br>CTTTGCCGATCATCCTTAGC | Grillet et al., 2018 |
| qRT_IMA1_Fw<br>qRT_IMA1_Rv | *IMA1* | TGATTGTAATTTAGGAGGAAACAAAA<br>TCAATCCACAAGTAAACATCTATGG | Grillet et al., 2018 |
| qRT_NAS4_Fw<br>qRT_NAS4_Rv | *NAS4* | ACGACCAACTCGTAAACAAG<br>GAGAGTGTCGACATCTTCAC | This work |
| bHLH39_At3g5698<br>bHLH39_At3g5698 | *bHLH39* | CGTGACCGTCGCAGGAAAATTAAC<br>TCGCAGGAATGCTTAGCTTCTTCG | This work |
| qRT_ZIF1_Fw<br>qRT_ZIF1_Rv | *ZIF1* | ACAGTTGGACCAGCTGGTGC<br>GCTACTCCGACCACTACTATCAC | This work |
| qRT_NRAMP1_Fw<br>qRT_NRAMP1_Rw | *NRAMP1* | TGCTCTCATCGGTGGTTCAG<br>CAAACGGGAGCTCAAAGGAT | Ihnatowicz et al., 2014 |
| qRT_CYP71B5_Fw<br>qRT_CYP71B5_Rv | *CYP71B5* | GACGAGAGCCAATTGTTGGT<br>GAGGTTGGTCTCCGATTCAA | This work |
| DTX1_601F<br>DTX1_856R | *DTX1* | TTGTTTGGTCTGGGATGTAATGG<br>CGAGTTTCGGGTTAGGGAGAAG | This work |
| qRT_MYB10_Fw<br>qRT_MYB10_Rv | *MYB10* | CGCTGGATTGATGAGATGCGGAA<br>TGAAGTTGCCTCGTTTGAGACCTG | This work |
| qPCR_MYB72_Fw<br>qPCR_MYB72_Rv | *MYB72* | GGACGTGAAGCGAGGCAACTTTAG<br>GAAGGACGCGATCTTTGACCACTT | This work |
| qPCR_BTSL2_Fw<br>qPCR_BTSL2_Rv | *BTSL2* | CGGGGCAGAATCCATCTTAT<br>GTTGCAACAAGGAGCAAGAAG | Rodríguez-Celma et al., 2019 |
| qRT_BGLU45_Fw<br>qRT_BGLU45_Rv | *BGLU45* | TTAGAAGCTTTACAAGCAGCAATG<br>CCACACAAAATAACCCTTCACA | This work |

(listed in order of appearance in the text)

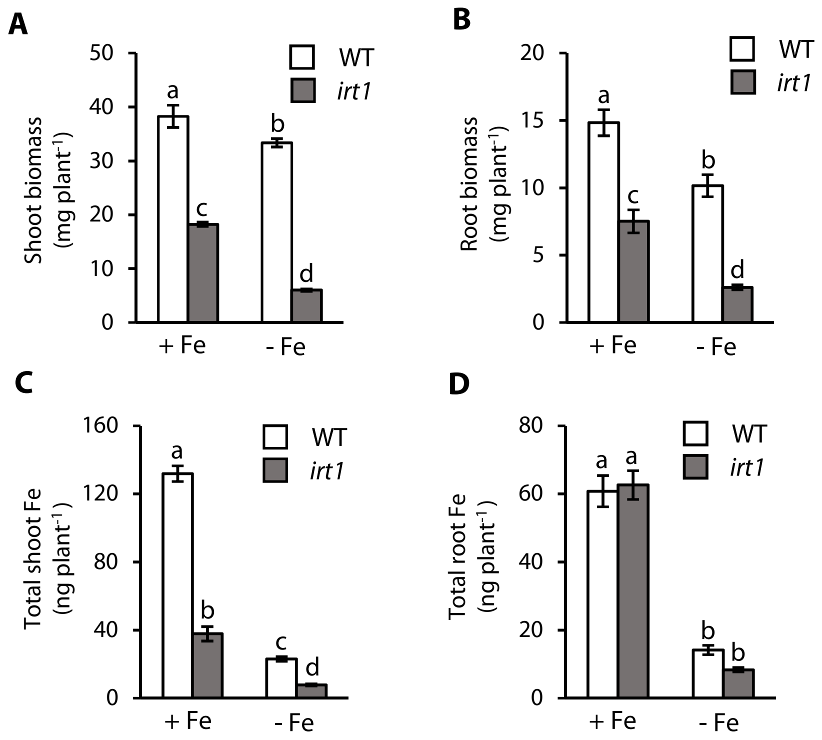


**Supplemental Figure S1. Details of the *irt1* phenotype. (A and B)** Fresh biomass of shoots **(A)** and roots **(B)** of 20-d-old WT and *irt1* seedlings grown in Fe-sufficient (+ Fe, 10 µM FeHBED) and Fe-deficient (- Fe, 0 µM FeHBED) agar-solidified 0.25x modified Hoagland's medium (EDTA-washed agar) for 10 d, after an initial cultivation in standard medium (5 µM FeHBED, unwashed agar) for 10 d, on vertically oriented petri plates. **(C and D)** Total Fe in shoots **(C)** and roots **(D)** of 20-d-old WT and *irt1* seedlings grown as in **(A)**. Bars represent arithmetic mean ± SD ($n$ = 3 pools of tissue from 15 (WT) or 30 (*irt1*) seedlings, with 15 seedlings cultivated per plate). Distinct letters indicate significant differences ($P < 0.05$, two-way ANOVA followed by Tukey´s HSD test). Data show one experiment representative of two **(A and B)** or three **(C and D)** independent experiments.

### Supplemental Figure S2

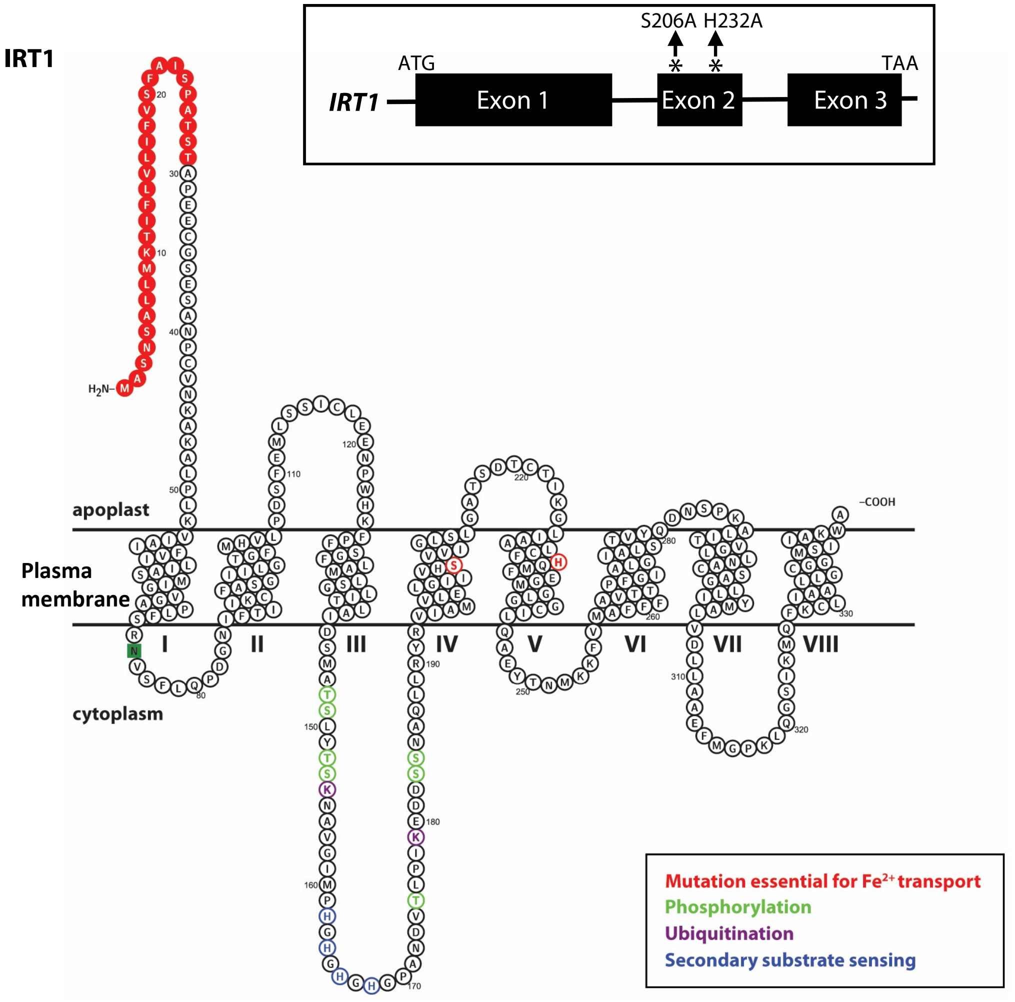


**Supplemental Figure S2. Topology model of IRT1 highlighting the amino acid residues mutated in this paper.** Shown are (i) the residues mutated in this paper (red), (ii) the predicted signal peptide (red circles), (iii) the cytosolic phosphorylation sites (green), (iv) cytosolic ubiquitination sites (purple) and (v) amino acids acting in secondary substrate sensing (navy blue). Box shows representation of the positions of nucleotides exchanged in the second exon of *IRT1* effecting alanine substitutions. Note that amino acid numbering is distinct from Rogers *et al.* (2000) because the *IRT1* cDNA employed in the earlier study was incomplete at the 5‘-end, leading to an N-terminal truncation of the IRT1 protein by eight amino acids.

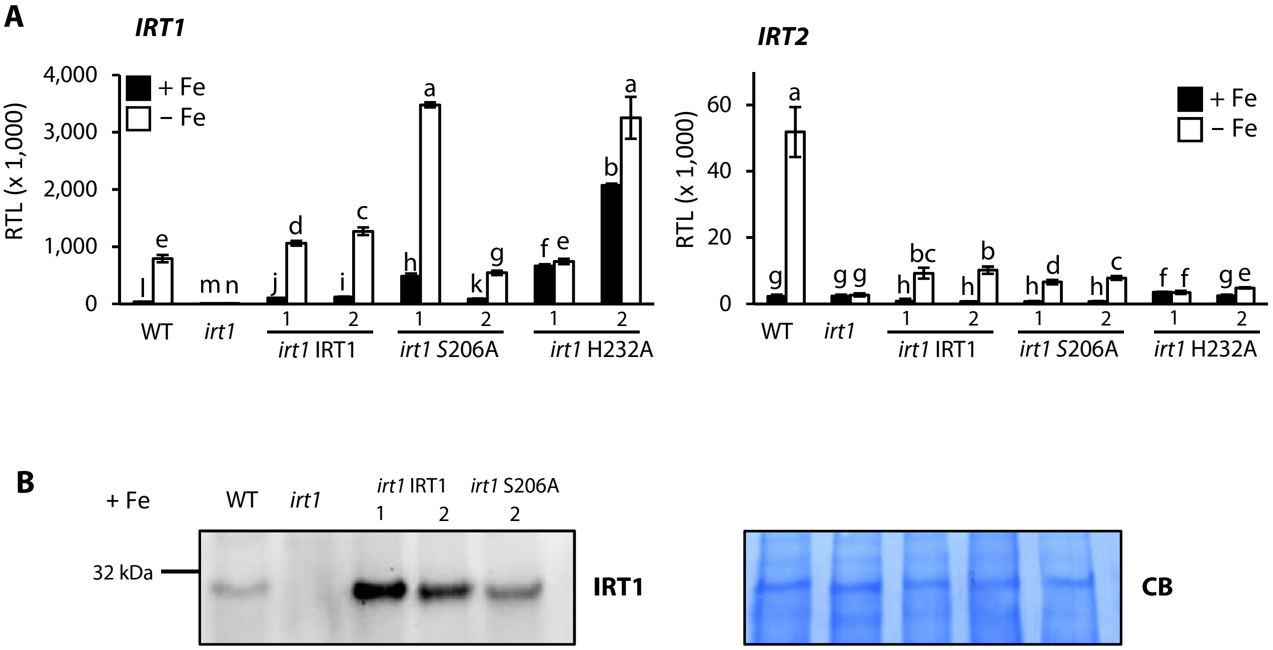


**Supplemental Figure S3. Details of *IRT1* expression in transport-inactive mutants. (A)** *IRT1* and *IRT2* relative transcript levels (RTL) in 15-d-old seedlings of WT, *irt1*, *irt1* IRT1, *irt1* S206A and *irt1* H232A lines. Seedlings were grown on Fe-sufficient (+ Fe, 10 µM FeHBED) and Fe-deficient (- Fe, 0 µM FeHBED) agar-solidified 0.25x modified Hoagland's medium (EDTA-washed agar) for 5 d, subsequent to an initial cultivation period of 10 d in standard medium (5 µM Fe-HBED; unwashed agar), on vertically oriented petri plates. Transcript levels are shown normalized to those of *UBQ10* and *EF1α*. **(B)** Immunoblot detection of IRT1 protein in 15-d-old seedlings of the WT, *irt1* and various transgenic lines. Total root protein extracts (20 µg) were separated on denaturing gels and blotted onto PVDF membranes. The Commassie Blue-stained (CB) PVDF membrane is shown as loading control. Seedlings were grown under + Fe conditions as described in **(A)**. Bars represent arithmetic mean ± SD ($n$ = 6 technical replicates using tissues pooled from 2 to 3 plates, each with 20 seedlings), and distinct letters indicate significant differences ($P < 0.05$) according to two-sample $t$-tests (*IRT1*) or Welch $t$-tests (*IRT2*) upon correction for multiple comparisons **(A)**. Data in **(A)** show one experiment representative of four independent experiments. Images in **(B)** are representative of two replicate membranes.

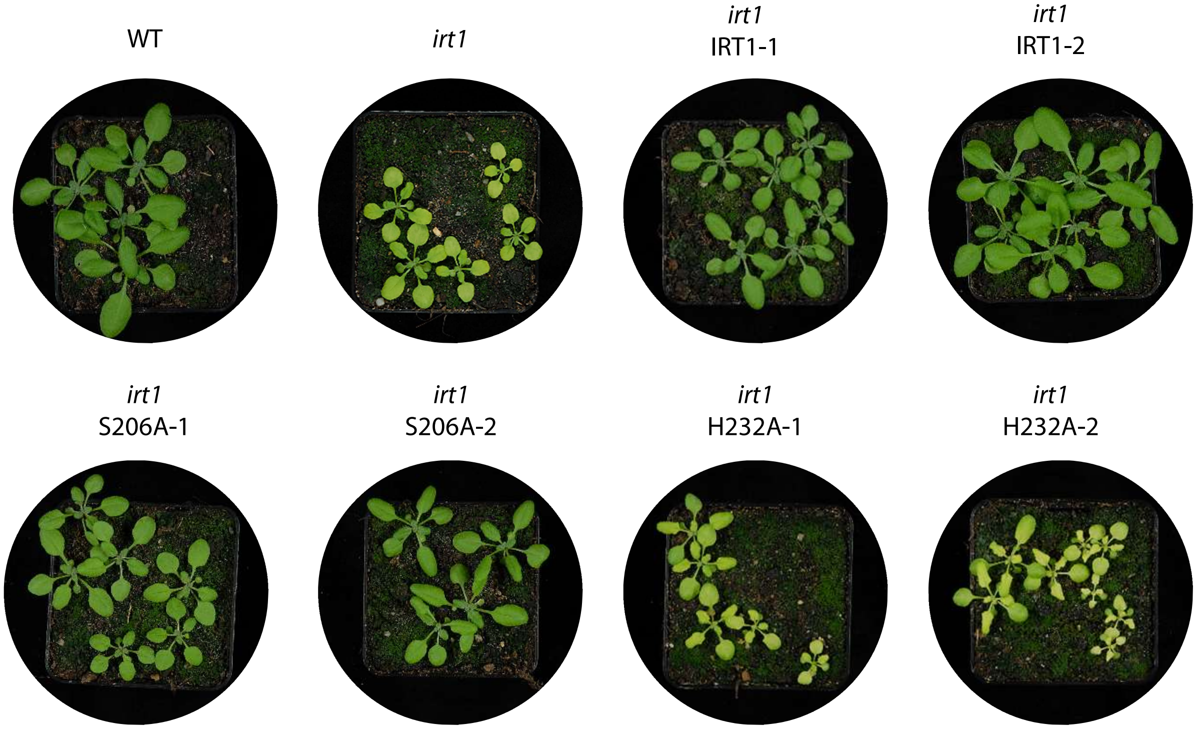


**Supplemental Figure S4. Photographs of Arabidopsis WT, *irt1*, *irt1* IRT1, *irt1* S206A, *irt1* H232A plants.** Shown are representative images of WT and *irt1* mutant plants as well as two independent transformant lines per construct upon cultivation on standard greenhouse soil for three weeks.

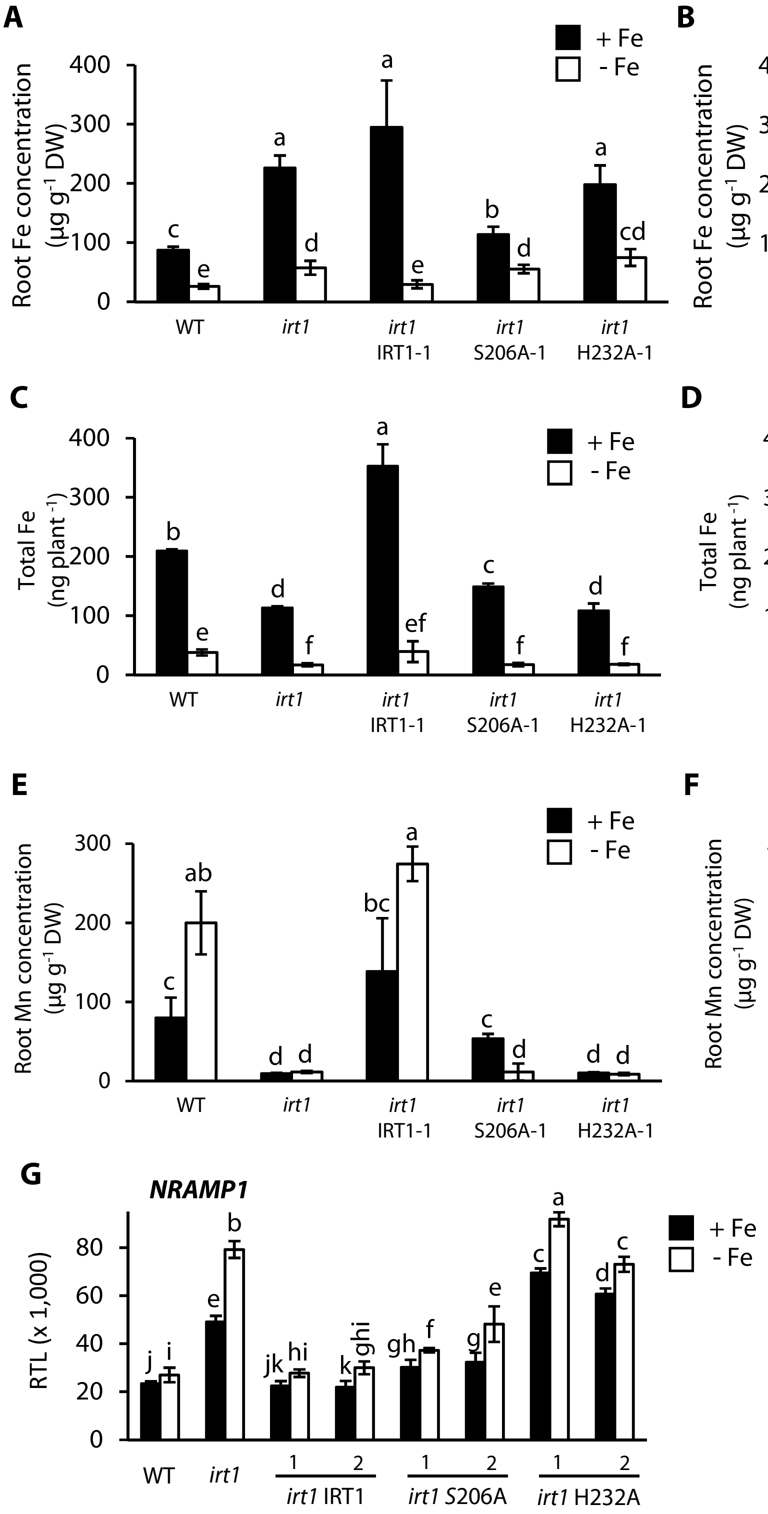


**Supplemental Figure S5. Root Fe concentration, total Fe per plant, root Mn concentration and *NRAMP1* expression. (A - F)** Root Fe concentrations **(A and B)**, total Fe per plant **(C and D)** and root Mn concentrations **(E and F)** in 20-d-old seedlings of WT, *irt1*, *irt1* IRT1, *irt1* S206A, *irt1* H232A grown on Fe-sufficient (+ Fe, 10 µM FeHBED) and Fe-deficient (- Fe, 0 µM FeHBED) agar-solidified 0.25x modified Hoagland's medium (EDTA-washed agar) for 10 d, subsequent to an initial cultivation period of 10 d in standard medium (5 µM FeHBED; unwashed agar) on vertically oriented petri plates. **(G)** *NRAMP1* relative transcript levels (RTL) in 15-d-old seedlings of WT, *irt1* and various transgenic *irt1* lines grown on + Fe and - Fe medium for 5 d. Transcript levels are shown normalized to those of *UBQ10* and *EF1α*. Bars represent arithmetic mean ± SD ($n$ = 3 pools of tissues from 15 to 30 seedlings, with 15 seedlings cultivated per plate **(A - F)**; $n$ = 3 technical replicates on cDNA obtained using tissue pooled from 2 to 3 plates, each with 20 seedlings **(G)**). Distinct letters indicate significant differences ($P < 0.05$) according to two-sample Student's $t$-tests upon correction for multiple comparisons. Data show one experiment representative of two **(A - F)** or four **(G)** independent experiments.

### Supplemental Figure S6

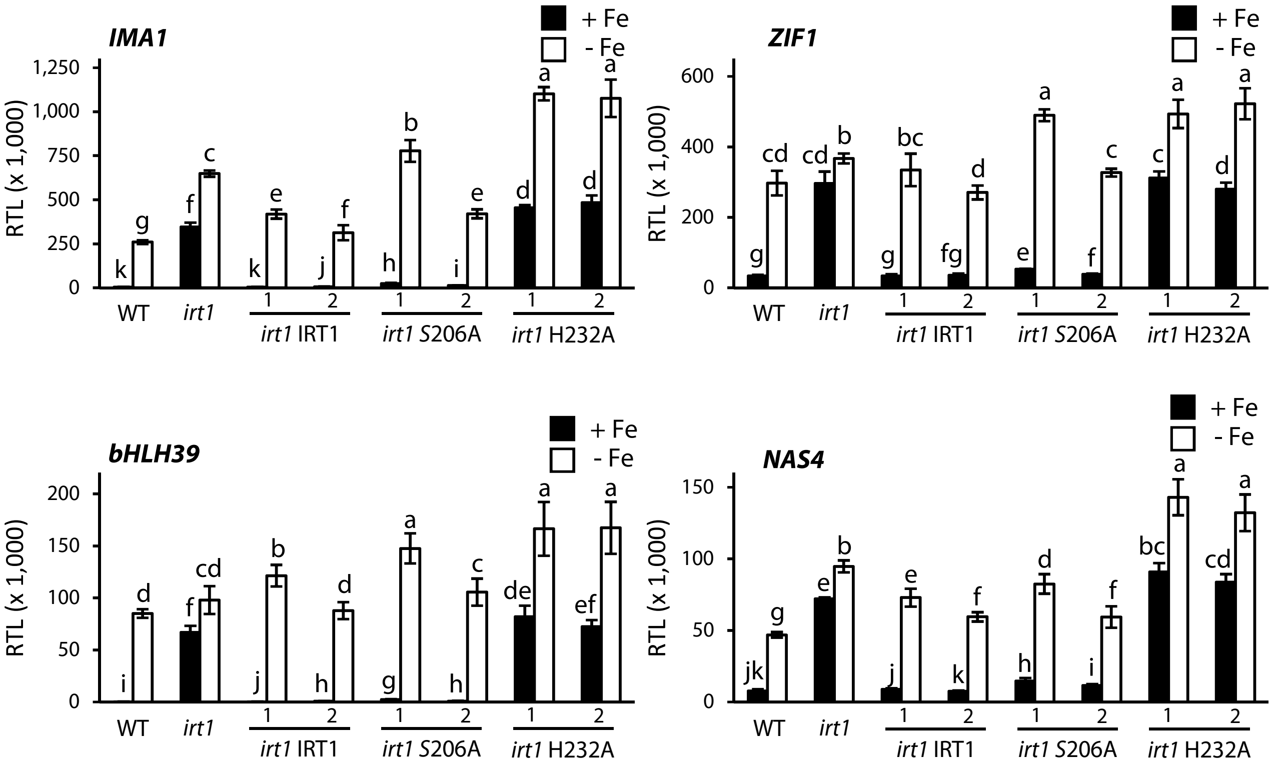


**Supplemental Figure S6. Fe deficiency responses are not constitutively activated in shoots of *irt1* S206A lines.** Relative transcript levels (RTL) of *IMA1*, *ZIF1*, *bHLH39* and *NAS4* in shoots of 15-d-old seedlings of the WT, *irt1*, *irt1* IRT1, *irt1* S206A, *irt1* H232A. RTL are shown normalized to both *UBQ10* and *EF1$\alpha$* as constitutively expressed control genes. Seedlings were grown on Fe-sufficient (+ Fe, 10 µM FeHBED) and Fe-deficient (- Fe, 0 µM FeHBED) agar-solidified 0.25x modified Hoagland's medium (EDTA-washed agar) for 5 d, subsequent to an initial cultivation period of 10 d in standard medium (5 µM FeHBED; unwashed agar), on vertically oriented petri plates. Bars represent arithmetic mean ± SD ($n$ = 6 technical replicates using tissue pooled from 2 to 3 plates, each with 20 seedlings). Distinct letters indicate significant differences ($P < 0.05$) according to two-sample Student's or Welch $t$-tests upon correction for multiple comparisons. Data show one experiment representative of three independent experiments.

### Supplemental Figure S7

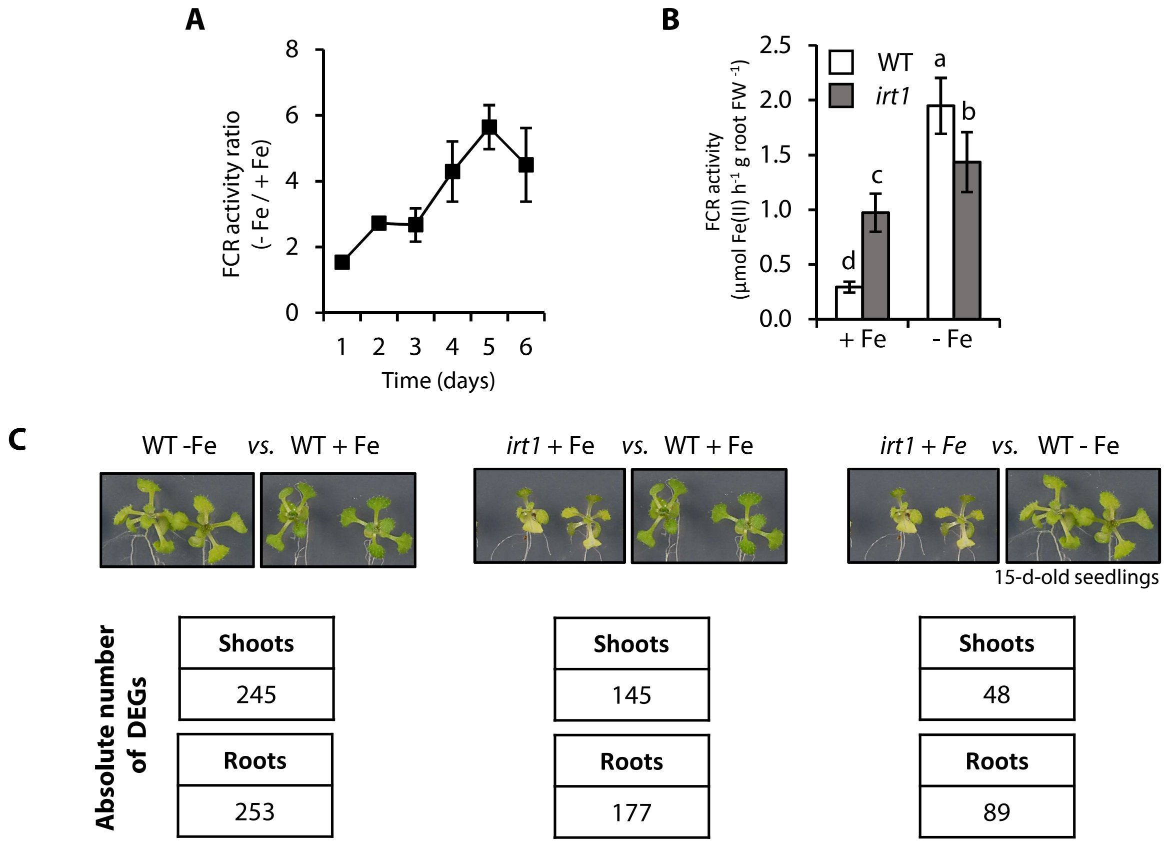


**Supplemental Figure S7. Experimental rationale of transcriptomic analysis. (A)** Time-course of root surface Ferric Chelate Reductase (FCR) activity in WT seedlings following transfer to Fe-deficient media. Seedlings were grown on Fe-sufficient (+ Fe, 10 µM FeHBED) and Fe-deficient (- Fe, 0 µM FeHBED) agar-solidified modified 0.25x Hoagland's medium (EDTA-washed agar) for 5 d, subsequent to an initial cultivation period of 10 d in standard medium (5 µM FeHBED; unwashed agar), on vertically oriented petri plates. Shown is the ratio between root surface FCR activity in - Fe and + Fe. **(B)** Root surface FCR activity in 15-d-old WT and *irt1* seedlings grown as in **(A)**. **(C)** Total numbers of differentially expressed genes (DEGs) in the microarray-based transcriptome analysis. Bars represent arithmetic mean ± SD ($n$ = 3 pools of 5 roots from 3 plates **(A)**; $n$ = 6 pools of 5 roots from plates **(B)**). Distinct letters indicate significant differences ($P < 0.05$) in two-way ANOVA followed by Tukey´s HSD test. Data in **(B)** show one experiment representative of three independent experiments.

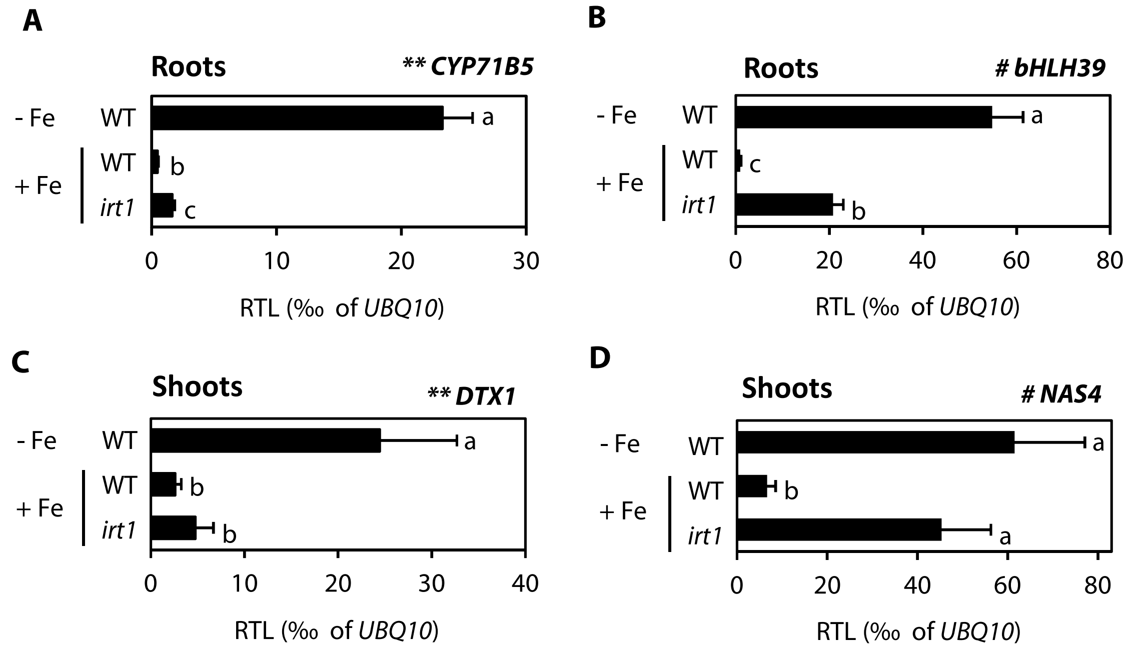


**E**

| **symbol** | **tairShortDescription** | **Fold Change in microarray** | | | | | |
|---|---|---|---|---|---|---|---|
| | | **-Fe *vs.* C (WT)** | **Direction** | ***irt1 vs.* WT (C)** | **Direction** | ***irt1* (C) *vs.* WT - Fe** | **Direction** |
| *CYP71B5* | cytochrome p450 71b5 | 23.3 | up | 1.7 | up | 13.5 | down |
| *bHLH39* | basic helix-loop-helix (bHLH) DNA-binding superfamily protein | 39.2 | up | 21.3 | up | 1.8 | down |
| *DTX1* | MATE efflux family protein | 3.7 | up | 1.27 | up | 2.9 | down |
| *NAS4* | nicotianamine synthase 4 | 7.6 | up | 5.3 | up | 1.4 | down |

**Supplemental Figure S8. Validation of microarray data using RT-qPCR. (A - D)** Relative transcript levels (RTL) of *NAS4, DTX1, bHLH39* and *CYP71B5* in 15-d-old WT and *irt1* seedlings grown on Fe-sufficient (+ Fe, 10 µM FeHBED) and Fe-deficient (- Fe, 0 µM FeHBED) agar-solidified 0.25x modified Hoagland's medium (EDTA-washed agar) for 5 d, subsequent to initial cultivation in standard medium (5 µM FeHBED; unwashed agar) for 10 d, on vertically oriented petri plates. Bars represent arithmetic mean ± SD ($n$ = 3 technical replicates on cDNAs obtained using tissues pooled from 2 to 3 plates, each with 20 seedlings). Distinct letters indicate significant differences ($P < 0.05$) in two-way ANOVA followed by Tukey´s HSD test (*NAS4, bHLH39* and *CYP71B5*) or two-sample Student's *t*-tests with correction for multiple comparisons (*DTX1*). **(E)** Table showing *CYP71B5, bHLH39*, *DTX1,* and *NAS4* expression fold change in microarray.

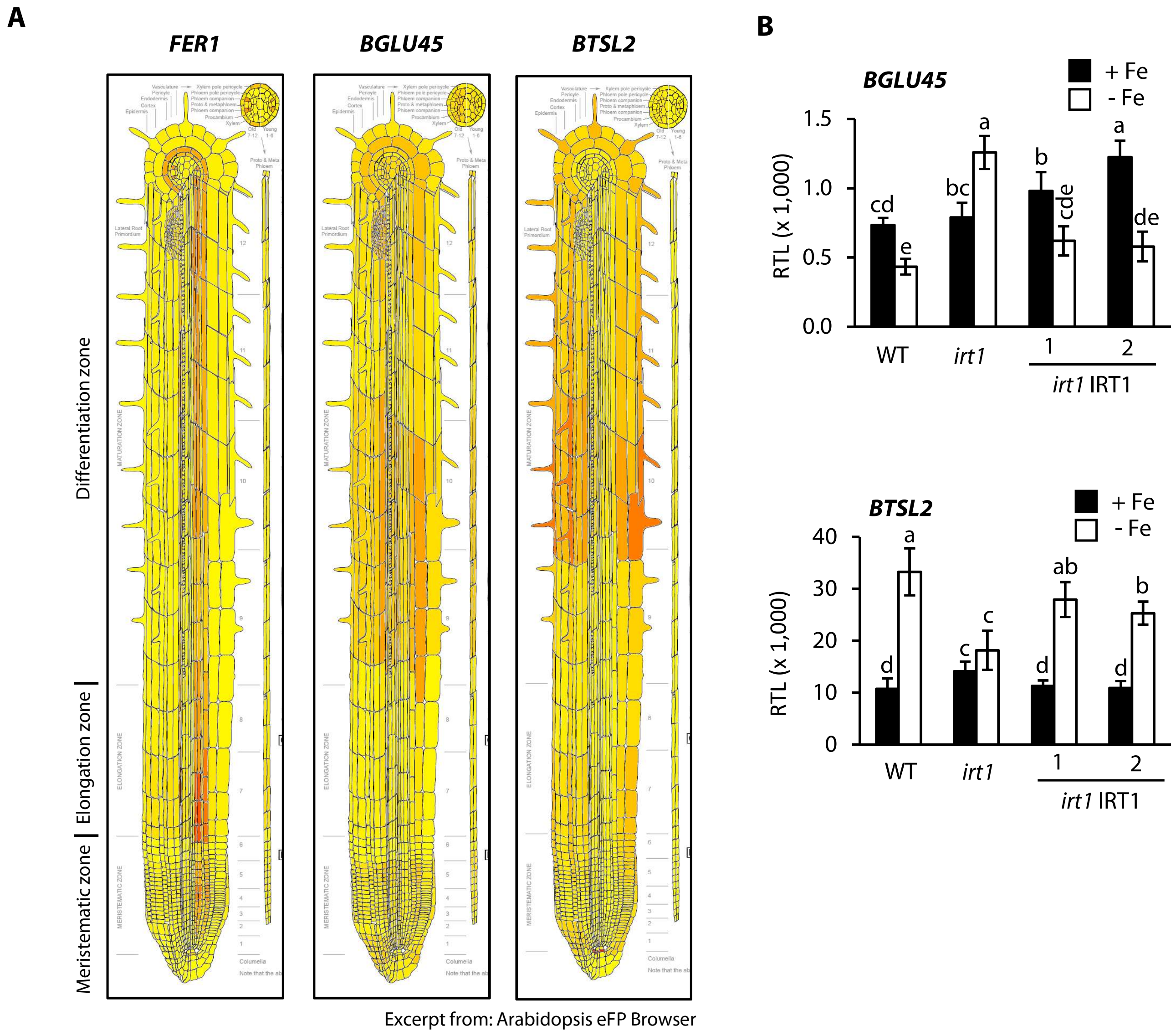


**Supplemental Figure S9. Lack of Fe-deficiency response in cortex-enriched transcripts in the *irt1* mutant. (A)** Cell-type specific expression in roots of 7-day-old seedlings according Brady et al (2007). **(B)** Relative transcript levels of *BGLU45* and *BTSL2* in shoots of 15-d-old seedlings of the WT, *irt1* and *irt1 IRT1*. RTL are shown normalized to both *UBQ10* and *EF1α* as constitutively expressed control genes. Seedlings were grown on Fe-sufficient (+ Fe, 10 µM FeHBED) and Fe-deficient (- Fe, 0 µM FeHBED) agar-solidified 0.25x modified Hoagland's medium (EDTA-washed agar) for 5 d, subsequent to an initial cultivation period of 10 d in standard medium (5 µM FeHBED; unwashed agar), on vertically oriented petri plates. Bars represent arithmetic mean ± SD ($n$ = 6 technical replicates on cDNA extracted using tissue pooled from 2 to 3 plates, each with 20 seedlings). Distinct letters indicate significant differences ($P < 0.05$) in two-way ANOVA followed by Tukey´s HSD test (*BGLU45*) or two-sample Student's $t$-test with correction for multiple comparisons (*BTSL2*). Data show one experiment representative of three independent experiments.

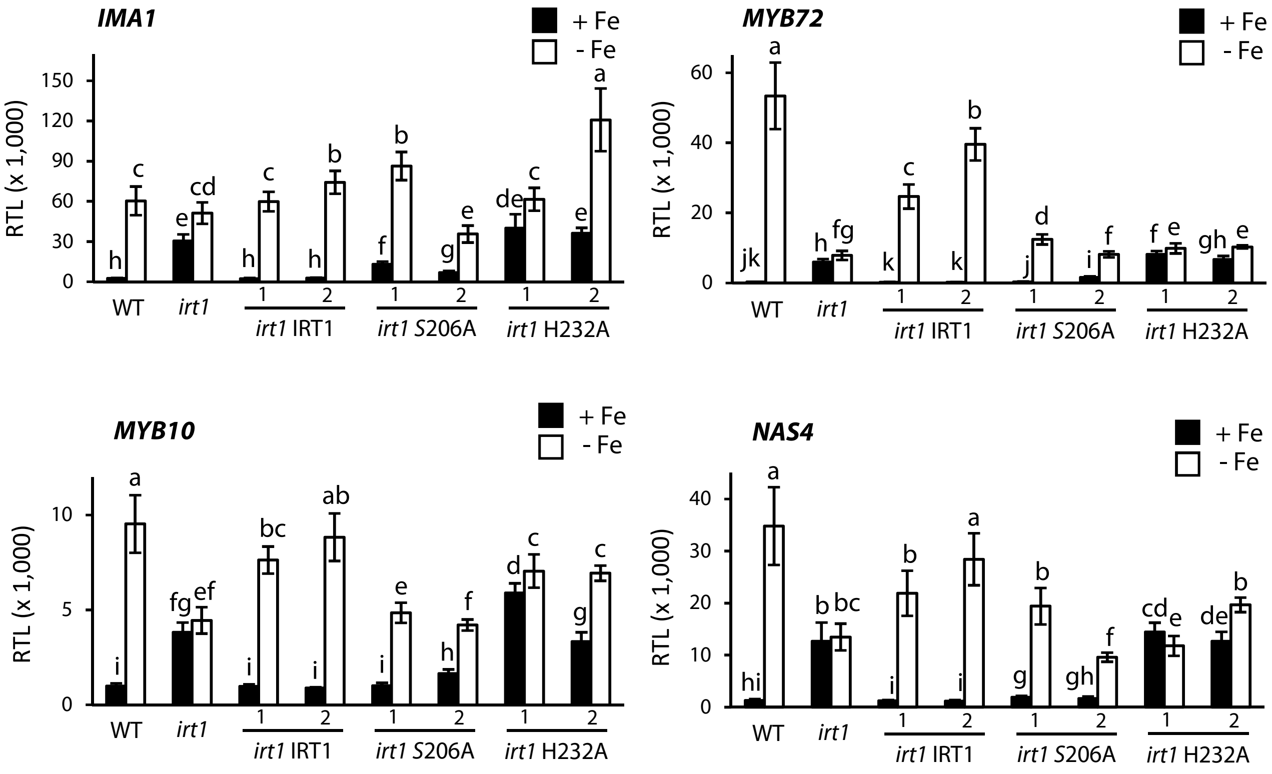


**Supplemental Figure S10. Transcript levels of candidate genes for roles in IRT1-dependent root-to-shoot translocation across genotypes.** Relative transcript levels (RTL) of *IMA1*, *MYB72*, *MYB10* and *NAS4* in roots of 15-d-old seedlings of WT, *irt1*, *irt1 IRT1, irt1* S206A*, irt1* H232A. RTL are shown normalized to both *UBQ10* and *EF1α* as constitutively expressed control genes. Seedlings were grown on Fe-sufficient (+ Fe, 10 µM FeHBED) and Fe-deficient (- Fe, 0 µM FeHBED) agar-solidified 0.25x modified Hoagland's medium (EDTA-washed agar) for 5 d, subsequent to an initial cultivation period of 10 d in standard medium (5 µM FeHBED; unwashed agar), on vertically oriented petri plates. Bars represent arithmetic mean ± SD ($n$ = 6 technical replicates on cDNA obtained using tissue pooled from 2 to 3 plates, each with 20 seedlings). Distinct letters indicate significant differences ($P$ < 0.05) according to two-sample Student's or Welch $t$-tests upon correction for multiple comparisons. Data show one experiment representative of three independent experiments.
